# Investigation of whether oxytocin and trust play a role in placebo effects of marketing actions

**DOI:** 10.1101/2022.12.01.518177

**Authors:** Daniela S. Schelski, Dirk Scheele, Liane Schmidt, René Hurlemann, Bernd Weber, Hilke Plassmann

**Affiliations:** Center for Economics and Neuroscience, University of Bonn, 53113, Bonn, Germany; Institute of Experimental Epileptology and Cognition Research, University of Bonn Medical Center, 53127, Bonn, Germany; Department of Social Neuroscience, Faculty of Psychology, Ruhr-University Bochum, 44780, Bochum, Germany; Control-Interoception-Attention Team, Institut du Cerveau et de la Moelle épinière, INSERM UMR 1127, CNRS UMR 7225, Sorbonne Université, 75013, Paris, France; Department of Psychiatry, School of Medicine & Health Sciences, University of Oldenburg, 26160, Bad Zwischenahn, Germany; INSEAD, 77300, Fontainebleau, France

**Keywords:** Placebo effects, Expectancy effects, Oxytocin, Trust, Marketing

## Abstract

Expectations about the quality of a medical treatment influence how much an inert treatment helps to improve patient well-being. Similarly, expectations about the quality of products influence how identical goods and services are evaluated differently after their consumption. One driver for such “placebo effects” in medical treatments is social cognition in the form of trust, which may be influenced by the hormone oxytocin. An open question is whether trust and oxytocin play similar roles in marketing placebo effects. To answer this question, we combined oxytocin administration (24 IU) and trust questionnaires in a pre-registered double-blind randomized between-subjects study design (*N_food tasting task_* = 223; *N_cognitive performance task_* = 202). We could not find evidence that oxytocin and trust play a role in placebo effects of marketing actions. Together with other recent null findings from oxytocin administration studies, these findings question the role trust might play in different types of placebo effects.

## Introduction

Expectations about the efficacy of a drug — or more generally the quality of a product — shape people’s experiences. Shapeable experiences can include subjective sensory perceptions such as pain, taste, and smell^1–5^, objective outcomes such as learning and cognitive performance^6–8^, and their underlying neurophysiological correlates^3,4,9–11^ and neuroendocrinological bases^12–14^, as well as subsequent motivated choices and behaviours^15,16^. Expectations can be generated based on signals from the environment. In the case of classic pain placebo effects, such environmental signals include being in a doctor’s office, seeing a syringe, verbal suggestions about the efficacy of treatment, and more implicit cues such as body language, facial expressions, and internal affective states^2^. In the case of more general everyday consumption decisions, price tags^8,17^, labels (e.g., indicating products to be low in calories^13^, organic^18^, or ethical)^19^, brand names^11,20^, and the mental representations created via advertisements^11,20–22^ have been identified as signals that generate quality expectations of economic goods, enhancing liking and consumption utility. Such expectancy effects have been termed “marketing placebo effects” (MPE)^8^.

The underlying neural pathways of pain placebo effects and marketing placebo effects are beginning to be understood^2,23–25^, but an interesting open question is why different signals have the ability to create expectations that colour experiences and motivated behaviours. One mechanism that has been suggested is that placebo effects on pain perceptions could be facilitated and increased by trust in the medical doctor, patient–doctor relationships characterized by warmth and competency^26^, and reduced social stress and anxiety^27^. A seminal study observed an increase in analgesic placebo response after intranasal administration of the neuropeptide oxytocin (OXT, 40 IU) in men. On this basis, the researchers proposed that the social relationship of patient and physician is essential for the formation of analgesic placebo effects^28^. Past research has shown that OXT has broad effects and plays a role in social attachment^29,30^, anxiety reduction^31^, and potentially social trust^32,33^, although its involvement in trust in humans has been debated more recently^34,35^. Follow-up research, however, could not confirm an effect of OXT on analgesic placebo responses, irrespective of participants’ sex, OXT dosage (24 or 40 IU), or placebo paradigm (verbal suggestion or conditioning)^27,36–38^.

These ideas have been parallelized by the importance of forming a relationship between customers and firms over time for the economic success of companies^39–41^: Trust in a certain brand, label, or specific marketer increases customers’ satisfaction and their loyalty.^42–46^ That is why companies spend a lot of their marketing budget on advertising, branding, and activities to improve customer relationships. However, it remains an open question whether trust in a marketer and its products actually increases MPE. Our research aimed at addressing this question by investigating the psychobiological link between trust in marketers and their actions and marketing placebo effects.

In this study, we tested whether OXT administration and reported trust in the marketer and their actions would increase MPE. Past consumer research provided first evidence that OXT modulated the consumer–brand relationship^47,48^ by increasing perceived brand competence and thus willingness to pay for branded products^47^. That is why we pre-registered the hypothesis that OXT would increase MPE by affecting the consumers’ trust in marketing actions. We also tested whether OXT increases MPE because trust in marketing actions increases consumers’ quality expectations.

To test our hypotheses, we administered 24 IU OXT to male participants who were randomly assigned to either a treatment group or a sham group in a double-blind procedure. Participants took part in two tasks previously designed to induce MPE (Figure 1): In a food tasting task participants (*N* = 223, *N*_sham_ = 111, *N*_OXT_ = 112) tasted identical foods with either a high or low price tag^3,4^ and an organic or neutral label^18^ and were then asked to rate how much they liked the taste. In a second task we explored the effect of an energy drink’s brand label on participants’ (*N* = 202, *N*_sham_ = 99, *N*_OXT_ = 103) cognitive performance in a numerical Stroop task^49^.

**Figure 1:**
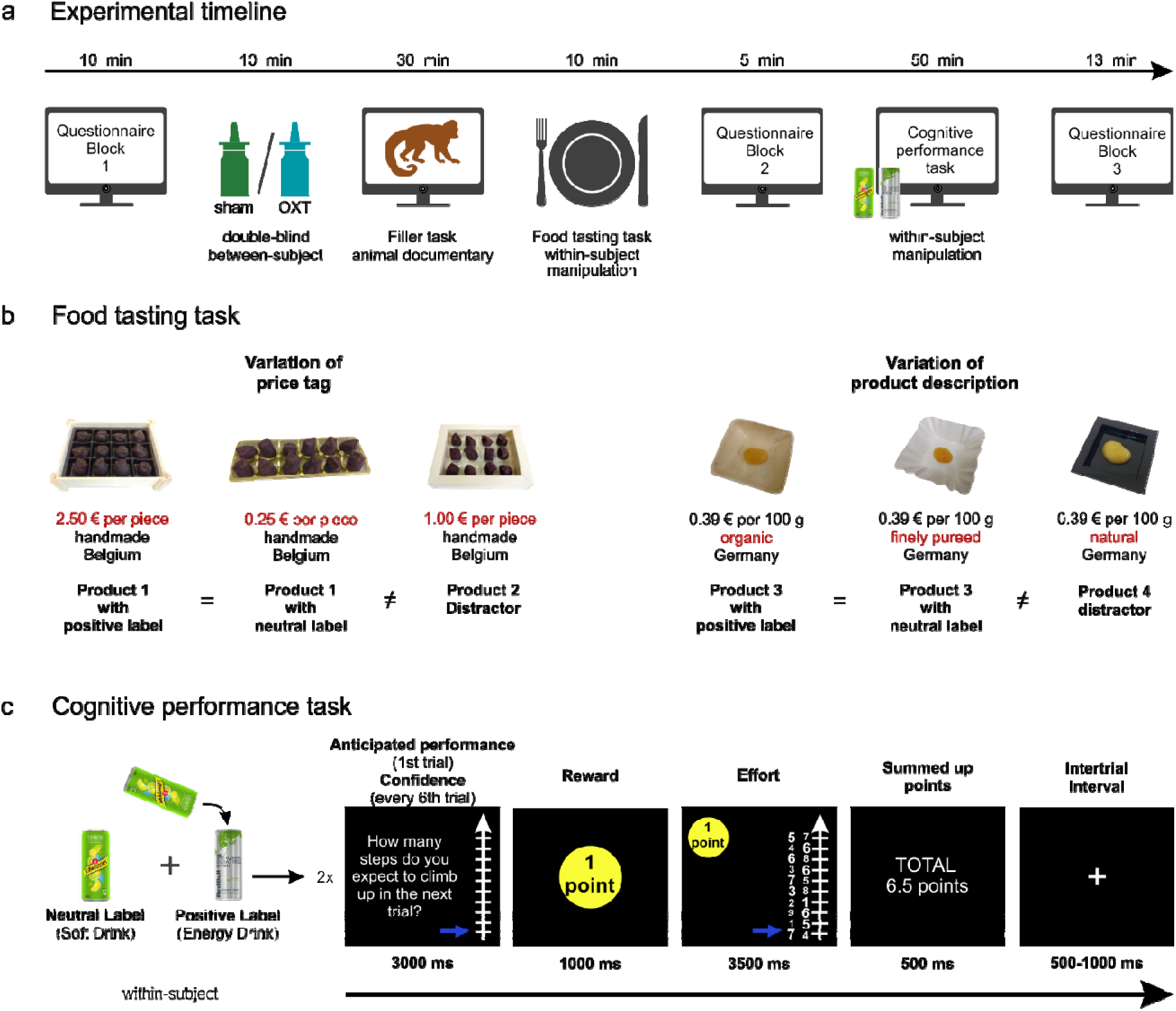
Overview of the experimental procedure with a food tasting task and a cognitive performance task. (**a**) After answering questionnaire block 1 (Autism Spectrum Quotient, general health questions, a general trust questionnaire, and Positive and Negative Affect Schedule), participants self-administered OXT or sham nasal spray. During a 30 min waiting period, participants watched an animal documentary as a filler task. Afterwards, in our main task, participants tasted different foods with different marketing labels and packaging (see **b**). In questionnaire block 2, we asked about trust in organic and expensive products. Afterwards, participants conducted a cognitive performance task with different brand labels (see **c**). Questionnaire block 3 comprised questions on trust in marketers of brand products, a manipulation check, and demographic and control questionnaires. (**b**) In the food tasting task, we varied within-participant prices of chocolates and labels of applesauce to trigger different expectations about the product’s taste. Information printed in red (for better visualization in this figure but not for the participants) is the manipulated marketing label of interest. The other information on the label was identical for all three products of each product type and served as distraction. (c) In a within-participant design, participants conducted the cognitive performance task twice, once after consuming a soft drink and once after consuming the same soft drink but labelled as Red Bull energy drink. Performance was measured for each trial during the effort phase in the form of the number of steps climbed up a ladder. Note that the question about the expected performance was used (1) as a measure of anticipated performance based on the response only before the first trial of each condition and (2) as a measure of confidence in performance every sixth trial. Abbreviations: OXT = oxytocin.

## Results

### Do marketing labels make identical food taste better and increase cognitive performance?

Conceptually replicating previous research, we found that marketing actions that increase expectations about how good a product might taste (i.e., high price tags, organic labels, and packaging) indeed let consumers enjoy products more. We found mixed results regarding how branding increased participants’ performance:

#### MPE effects on taste experiences

The food tasting task was designed to capture MPE on participants’ subjective taste experiences. We used either expensive or organic labels together with respective packaging and compared them to tasting identical products with neutral labels and more neutral packaging (labels and packaging jointly described as “labels” in the following). With this manipulation we aimed at generating higher versus lower expectations about how good the food would taste^3,4,18^. Participants rated their subjectively experienced taste pleasantness on a 9-point Likert scale. The labels inducing positive taste expectations led to a statistically significant increase in taste pleasantness by 5.3% (mean ± SD neutral labels: 6.17 ± 0.78, mean ± SD positive labels: 6.50 ± 0.78, *β* = 0.34, SE = 0.10, 95% CI [0.15, 0.53], *t* = 3.49, *p* < 0.001, *d* = 0.44; see Supplementary Table S1 and Figure 2a).

**Figure 2:**
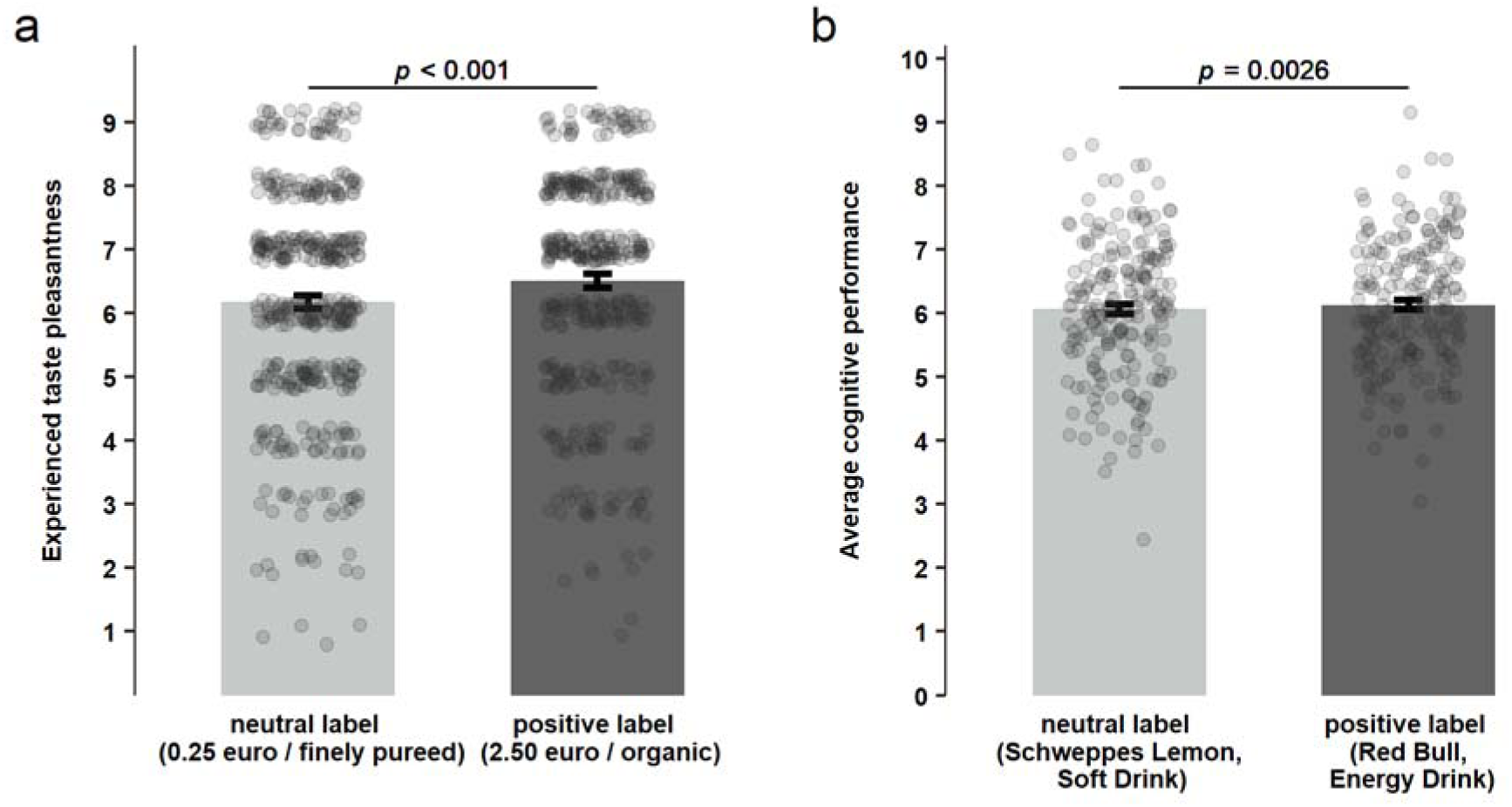
Effects of expectation enhancing (i.e., positive) marketing actions on experienced taste pleasantness and cognitive performance. (**a**) Positive labels significantly increased taste pleasantness ratings compared to neutral labels across both product types (*β* = 0.34, SE = 0.10, 95% CI [0.15, 0.53], *t* = 3.49, *p* < 0.001, *d* = 0.44). (**b**) Positive (energy drink) labels significantly enhanced the average cognitive performance across all trials compared to neutral (soft drink) labels (*β* = 0.06, SE = 0.02, 95% CI [0.02, 0.10], *t* = 3.02, *p* = 0.003). Note that both label effects, on experienced taste pleasantness and cognitive performance, are shown across all participants — sham and OXT group. It is important to note that the main effect on cognitive performance (**b**) was driven by a significant interaction as described below. Individual observations (**a**: *N* = 446, two observations per participant, one for each product; **b**: *N* = 202, one average cognitive performance per participant) are shown with grey dots. Error bars denote 95% confidence interval of the means.

A post hoc separate paired and two-sided *t*-test for both labels with Holm correction for multiple testing revealed a statistically significant influence of the price label on taste pleasantness (mean ± SD low price tag: 6.65 ± 1.64, mean ± SD high price tag: 7.10 ± 1.46, *t*(222) = 4.18, *p*_corr._ < 0.001, *d* = 0.28, 95% CI of the difference [0.24, 0.67]; see Supplementary Figure S1a). Taste pleasantness increased by 6.8% due to the higher price tag. However, the organic label led only to a marginal, statistically nonsignificant increase in taste pleasantness of 3.7% (mean ± SD neutral label: 5.70 ± 1.70, mean ± SD positive label: 5.91 ± 1.76, *t*(222) = 1.84, *p*_corr._ = 0.068, *d* = 0.12, 95% CI of the difference [-0.02, 0.44]; see Supplementary Figure S 1b). Since we were more generally interested in the psychobiological mechanisms of marketing placebo effects induced by various marketing actions, we pooled both product types for all further analysis.

#### MPE effects on cognitive performance

The second task was designed to capture MPE on participants’ motivated behaviours — in this case, performance in a numerical Stroop task under time pressure. We measured participants’ cognitive performance under two conditions: once after consuming a soft drink and once while believing they had consumed an energy drink from a well-known brand. Unbeknownst to the participants, the drink delivered in the energy drink can and advertised for its effects on cognitive performance was identical to the soft drink. We captured cognitive performance by the number of correct answers in a numerical Stroop task. In this task participants had to indicate which number of a pair was numerically greater (Figure 1c). To increase the difficulty, we varied the physical size of the numbers to be congruent or not with the numerical size. In a robust linear mixed-effects model, we found a statistically significant main effect of label across both treatment groups (sham and OXT) (*β* = 0.06, SE = 0.02, 95% CI [0.02, 0.10], *t* = 3.02, *p* = 0.003; Figure 2b, Supplementary Table S2). However, this main effect was driven by the OXT treatment group, as reported below.

To conclude, we did replicate previous findings that marketing actions such as price tags, production labels, and packaging significantly increased subjective enjoyment of products. Our main question of interest was whether this might be due to a social relationship between consumers and marketers. We tested this question in two ways: (1) by testing whether OXT would increase MPE and (2) by testing whether reported trust in marketing actions would increase MPE.

### Does OXT administration increase MPE?

#### Effects of OXT for MPE during subjective taste experiences

To test our hypothesis that OXT can boost MPE on subjective liking of food, we calculated an MPE score for each participant (MPE score = taste pleasantness rating for positive label minus taste pleasantness rating for neutral label). The MPE score is a measure of the strength of the marketing action’s effect on each individual’s liking of the food’s taste.

Contrary to our hypothesis, we did not find any statistically significant influence of OXT on the MPE (mean ± SD sham group: 0.29 ± 1.13, mean ± SD OXT group: 0.38 ± 1.09, *β* = 0.09, SE = 0.16, 95% CI [-0.22, 0.40], *t* = 0.55, *p* = 0.58, *d* = 0.08; Figure 3a and Supplementary Table S3). The influence of OXT on the MPE score did not differ statistically significantly for the two product types, as indicated by the interaction term (*β* = −0.09, SE = 0.31, 95% CI [-0.70, 0.53], *t* = −0.28, *p* = 0.78).

**Figure 3:**
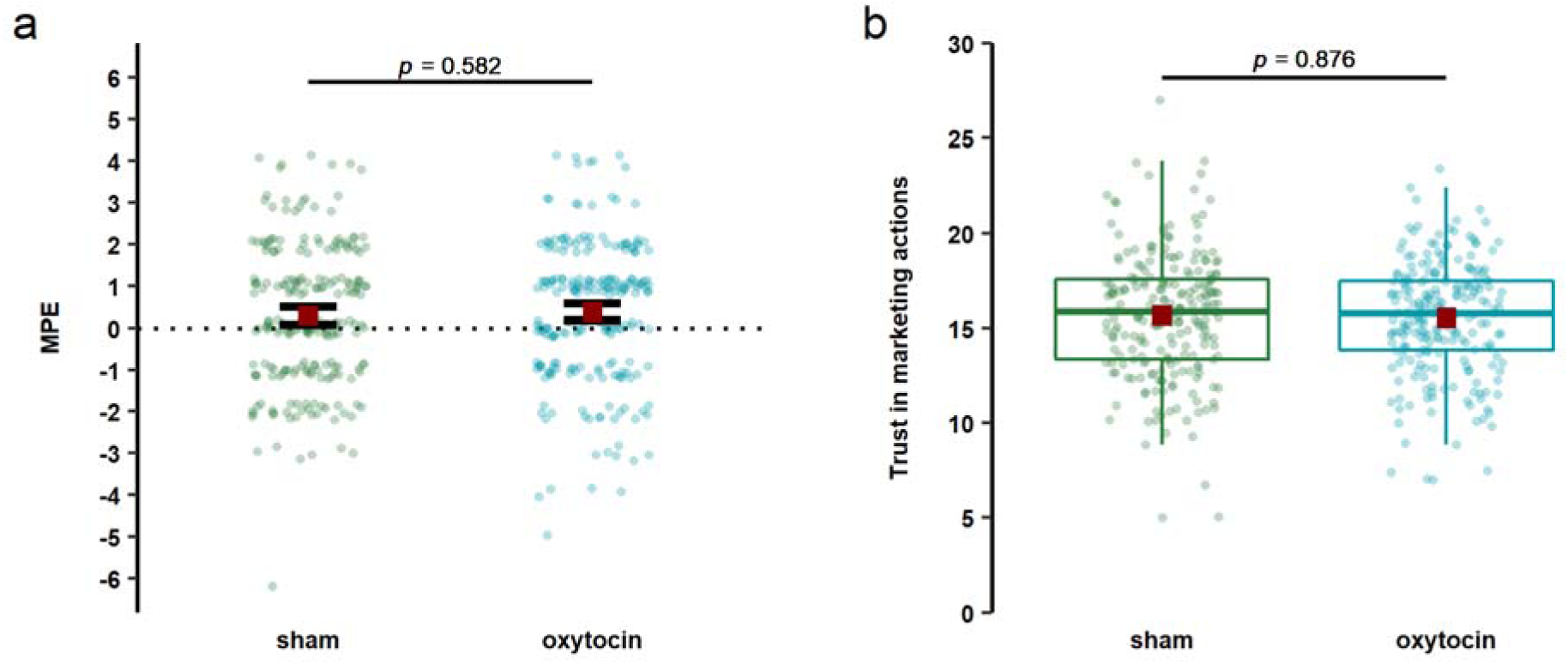
Evaluation of OXT’s impact on MPE on experienced taste pleasantness and OXT’s impact on trust in marketing actions. OXT did not significantly alter (**a**) MPE (*β* = 0.09, SE = 0.16, 95% CI [-0.22, 0.40], *t* = 0.55, *p* = 0.58, *d* = 0.08) or (**b**) trust in marketing actions (*β* = −0.05, SE = 0.33, 95% CI [-0.71, 0.60], *t* = - 0.16, *p* = 0.88, *d* = 0.02). Dots represent individual data points of participants for both products pooled (*N*_sham_ = 222, *N*_OXT_ = 224). Red squares show mean values. We depicted mean values with error bars for discrete Likert-scale data and used boxplots for continuous data. Error bars for discrete Likert-scale data (**a**) denote 95% confidence intervals and whiskers of boxplot for continuous data and (**b**) denote 1.5 x interquartile range, with the black line representing the median value. MPE = marketing placebo effect; OXT = oxytocin.

The null effect of treatment across both product types was robust when controlling for the change in positive and negative affect from before to after the experiment, general preference for chocolate truffles and applesauce, treatment guess, age, income, autism spectrum quotient, and general trust (see model 2 in Supplementary Table S3).

In order to interpret the null finding of our main hypothesis, we quantified the evidence in favour of an OXT effect on MPE with a Bayesian analysis. The Bayes factor BF10 of 0.018 indicates that the null model is 55 times more likely than the alternative model and thus supports the null effect with very strong evidence^50^. Put differently, the Bayesian analysis supports the null finding from the frequentist statistical linear mixed-effects model showing that there is a null effect of OXT on MPE. Adjusting the default prior (JZS prior with an *r* scale of 0.5) of the treatment regressor did not substantially change the evidence in favour of the null model (see Supplementary Table S4).

#### Effects of OXT for MPE during cognitive performance

In a robust linear mixed-effects model, we estimated how drink label, treatment, and several other regressors accounting for the features of the numerical Stroop task (i.e., task difficulty, reward level, run, drink order, trial number), interaction of treatment and label, and other possible interactions (interaction of treatment and difficulty, interaction of treatment and reward) influenced cognitive performance on a single trial level with random intercepts for participants.

OXT treatment did not statistically significantly influence cognitive performance (*β* = 0.02, SE = 0.14, 95% CI [-0.26, 0.30], *t* = 0.14, *p* = 0.89). We found a statistically significant main effect of run (*β* = 0.97, SE = 0.04, 95% CI [0.89, 1.05], *t* = 24.39, *p* < 0.001), indicating a better performance in the second run (probably due to learning effects). We also found a statistically significantly negative main effect of trial (*β* = −0.01, SE = 0.0007, 95% CI [-0.007, - 0.004], *t* = −7.99, *p* < 0.001), indicating that performance got worse in later trials (probably due to fatigue effects within a run). Moreover, we found a statistically significant positive main effect of reward (*β* = 0.06, SE = 0.02, 95% CI [0.02, 0.10], *t* = 3.15, *p* = 0.002), probably due to a higher motivation to perform better when more money was at stake. Trial difficulty had a statistically significant main effect on performance (*β* = −2.34, SE = 0.02, 95% CI [-2.38, −2.30], *t* = −117.62, *p* < 0.001) such that performance was lower in difficult trials.

A statistically significant main effect of label (*β* = 0.06, SE = 0.02, 95% CI [0.02, 0.10], *t* = 3.02, *p* = 0.003) indicated that across both treatment groups (sham and OXT) performance was higher in the energy drink condition than in the soft drink condition. Notably, drink label statistically significantly interacted with the treatment such that the OXT group showed a stronger label effect than the sham group (*β* = 0.09, SE = 0.04, 95% CI [0.02, 0.17], *t* = 2.38, *p* = 0.017; Figure 4a and Supplementary Table S2). More specifically, the average cognitive performance of the sham group did not change due to the energy drink label (mean ± SD performance with soft drink label: 6.06 ± 1.06; mean ± SD performance with energy drink label: 6.06 ± 0.94), while the performance of the OXT group slightly increased (mean ± SD performance with soft drink label: 6.06 ± 1.03; mean ± SD performance with energy drink label: 6.18 ± 1.04). The interaction effect of treatment and drink label remained statistically significant after controlling for personality, demographic variables, and changes in mood states (see model 2 in Supplementary Table S2).

**Figure 4:**
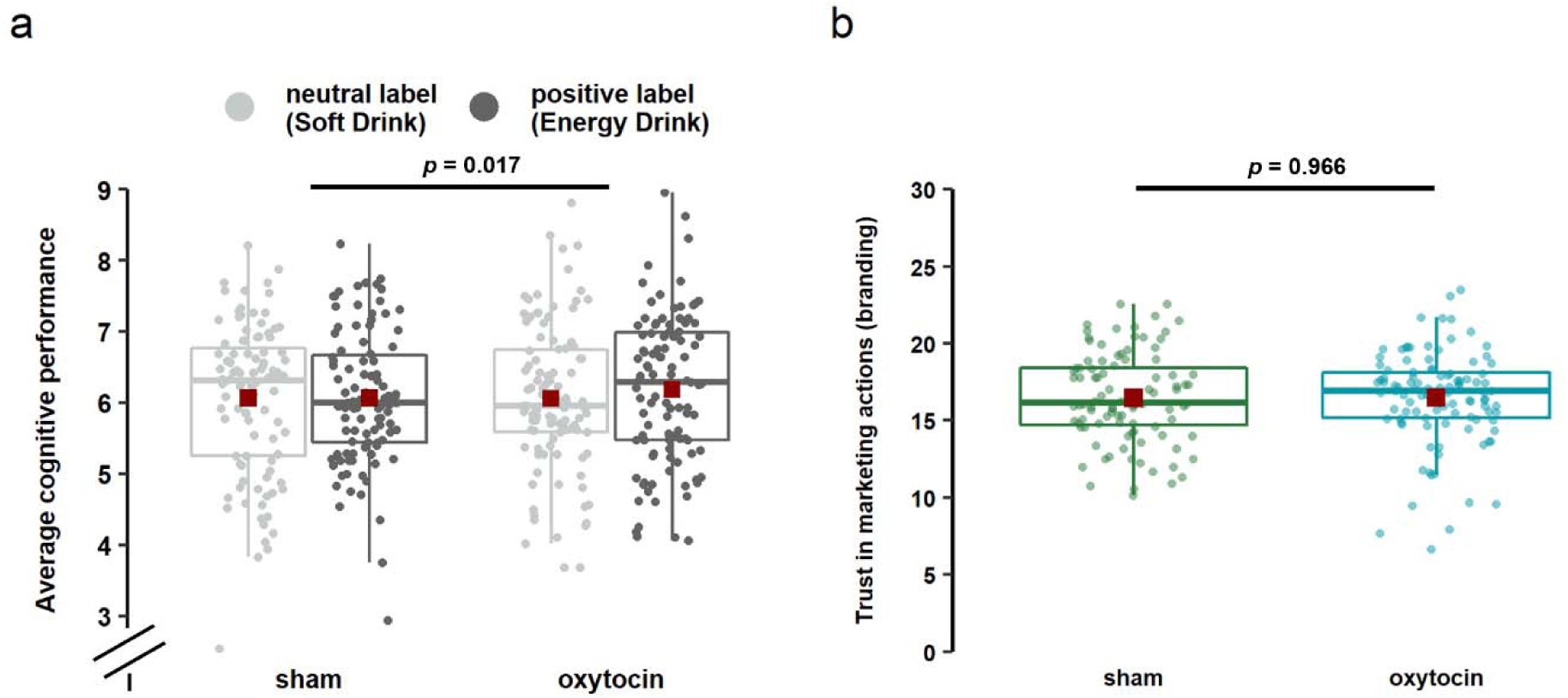
Evaluation of OXT impact on MPE on cognitive performance and OXT impact on trust in marketing actions. (**a**) OXT treatment significantly increased the MPE effect on the cognitive performance in the numerical Stroop task (*β* = 0.09, SE = 0.04, 95% CI [0.02, 0.17], *t* = 2.38, *p* = 0.017). Performance was measured within-participant in two runs of the numerical Stroop task, one for the soft drink condition and one for the energy drink condition. Note that the significant interaction of label and OXT treatment in the frequentist analysis could not be confirmed with a Bayesian analysis. (**b**) Oxytocin had no significant effect on trust in marketing actions (i.e, branding) (*t*(200) = −0.04, *p* = 0.966, *d* = 0.006, 95% CI of the difference [-0.85, 0.81]). Dots represent individual data points of participants (*N*_sham_ = 99, *N*_OXT_ = 103). Red squares show mean values. We used boxplots for continuous data. Whiskers of boxplots denote 1.5 x interquartile range; black lines represent the median value. MPE = marketing placebo effect; OXT = oxytocin.

To test that this interaction was not a false positive finding^51^, we additionally quantified the evidence in favour of this interaction effect by applying Bayesian statistics. Note that this Bayesian analysis was not pre-registered in advance. The Bayes factor BF10 of 0.16 indicates that the null model is 6.25 times more likely than the alternative model and thus supports a null effect with substantial evidence^50^. Hence, the Bayesian analysis disagrees with our frequentist statistical linear mixed-effects model result, as it does not provide evidence in favour of the interaction of OXT treatment with drink label. Adjusting the default prior (JZS prior with an *r* scale of 0.5) of the treatment regressor did not change the evidence in favour of the null model (see Supplementary Table S5).

To conclude, our results seem to suggest that OXT administration does not reliably increase MPE across different marketing actions and across different tasks (i.e., judging taste pleasantness and performing a cognitive task).

### Does reported trust in marketing actions increase MPE, and is it linked to OXT?

Because OXT’s role in human trust in an economic context has been debated recently^34,35^, we next investigated (1) whether reported trust in our employed marketing actions would increase MPE and (2) whether OXT would increase such feelings of trust in marketing actions.

#### Effects of reported trust in marketing actions for MPE during subjective taste experiences

To test this idea, we examined the link between trust in organic labels and high price tags and MPE. Higher levels of reported trust in organic labels and price tags did not show a statistically significant link to higher MPE (*β* = 0.09, SE = 0.09, 95% CI [-0.08, 0.26], *t* = 1.07, *p* = 0.28; Supplementary Figure 2a and Supplementary Table S6).

We further conducted an exploratory and not pre-registered analysis that tested whether the magnitude of MPE could be linked to label-induced (i.e., expensive and organic labels) quality and taste expectations. We found that only higher label-induced taste expectations (*β* = 0.20, SE = 0.10, 95% CI [0.008, 0.39], *t* = 2.05, *p* = 0.041) but not quality expectations (*β* = −0.13, SE = 0.10, 95% CI [-0.32, 0.06], *t* = −1.30, *p* = 0.195) significantly correlated with an increased MPE (see Supplementary Figure 2b and Supplementary Table S7).

Additional explorative analyses revealed a significant correlation of trust in marketing actions with quality expectations (*β* = 0.73, SE = 0.05, 95% CI [0.63, 0.83], *t* = 13.85, *p* < 0.001) and taste expectations of expensive and organic products (*β* = 0.70, SE = 0.06, 95% CI [0.57, 0.83], *t* = 10.85, *p* = < 0.001) (see Supplementary Table S8).

#### Effects of reported trust in marketing actions for MPE during cognitive performance

We explored whether trust in branding and anticipated cognitive performance due to branding (i.e., expected performance measured before the first trial of the first run) moderated the label effect on actual average performance (see Supplementary Table S9). Neither trust in branding (*β* = 0.09, SE = 0.13, 95% CI [-0.17, 0.35], *t* = 0.66, *p* = 0.510) nor anticipated performance (*β* = −0.08, SE = 0.13, 95% CI [-0.34, 0.18], *t* = −0.61, *p* = 0.546) significantly moderated the effect of the drink label on cognitive performance. Thus, MPE on cognitive performance did not significantly rely on trust in or expectations of a branded energy drink. Note also that trust in branding and anticipated performance in the first run did not significantly correlate (*r*(200) = −0.028, *p* = 0.692, Pearson correlation). These findings might explain why we did not find a label effect on cognitive performance in the sham group.

In two further linear models (Supplementary Tables S10 and S11) we explored whether anticipated performance and confidence in performance in the first run was dependent on drink label. In the sham group, drink label did not statistically significantly affect anticipated performance (*β* = 0.30, SE = 0.31, 95% CI [-0.31, 0.91], *t* = 0.98, *p* = 0.331) or confidence in performance (*β* = 0.16, SE = 0.19, 95% CI [-0.22, 0.53], *t* = 0.82, *p* = 0.411).

Moreover, in the same linear models (Supplementary Tables S10 and S11) we tested whether OXT treatment or trust in branding moderated the label effect on anticipated performance and confidence in performance. In other words, we wanted to know whether higher trust in branding and/or OXT treatment related to higher label-induced expectations of the drink. We did not find a statistically significant moderation of the effect of drink label on anticipated performance (*β* = 0.20, SE = 0.62, 95% CI [-1.02, 1.43], *t* = 0.33, *p* = 0.741) or confidence in performance (*β* = −0.28, SE = 0.38, 95% CI [-1.03, 0.46], *t* = −0.75, *p* = 0.453) by OXT treatment. Moreover, trust in branding did not statistically significantly moderate the effect of drink label on anticipated performance (*β* = −0.12, SE = 0.32, 95% CI [-0.74, 0.51], *t* = −0.37, *p* = 0.709) or confidence in performance (*β* = −0.04, SE = 0.19, 95% CI [-0.42, 0.34], *t* = −0.21, *p* = 0.836).

#### Reported trust in marketing actions and OXT

Last, we tested whether OXT increased reported measures of trust in marketing actions. Our analysis revealed no statistically significant difference in the reported trust in marketing actions used in the tasting task for the sham and OXT groups (mean ± SD sham group: 15.66 ± 2.70, mean ± SD OXT group: 15.52 ± 2.40, *β* = −0.05, SE = 0.33, 95% CI [-0.71, 0.60], *t* = −0.16, *p* = 0.88, *d* = 0.02; see Figure 3b and Supplementary Table S12). This null effect of OXT on trust in marketers remained robust when controlling for various personality and demographic variables (see model 2 in Supplementary Table S12). We also did not find a significant difference in brand trust for the sham and OXT groups for the cognitive performance task (mean ± SD in sham group: 16.46 ± 2.9, mean ± SD in OXT group: 16.48 ± 3.06, *t*(200) = −0.04, *p* = 0.966, *d* = 0.006, 95% CI of the difference [-0.85, 0.81] unpaired, two-sided *t*-test; see Figure 4b).

## Discussion

In this study, we investigated whether social cognition in the form of trust in marketers and their activities (such as how they label, package, and position their products) and OXT are drivers of why marketing actions change consumers’ experiences, even if the products themselves are unchanged. We replicated previous findings that positive marketing actions (i.e., those that increase positive expectations about the product) change the subjective consumption experience but not subsequent, more objective behaviour such as cognitive performance. However, we did not find evidence that OXT and through it trust^28^ could boost such expectancy effects.

Because OXT’s role in human trust in an economic context has been debated more recently^34,35^, we examined whether reported trust in our employed marketing actions instead of OXT per se would increase MPE. Although we observed that trust in the different marketing actions increased taste (but not performance) expectations, we did not find evidence that higher levels of reported trust are linked to higher MPE.

Last, to address the question of whether OXT administration is linked to trust in our setting at all, we also tested whether there is a link between OXT administration and reported trust in marketing actions, but again we could not find a significant relationship between these constructs.

Taken together, the lack of evidence of an OXT effect on placebo effects on subjective experiences is in line with more recent research, which has not replicated the initially observed positive effect of OXT on subjective experiences such as analgesic placebo responses^27,28,36–38^. Our findings provide important support for these other findings because one strength of our paper is its statistical power to detect even subtle effects due to the large sample size of 223 participants (i.e., 111–112 participants per experimental cell, which is about three times the sample size per cell of previous studies). However, as we did not test the effects of OXT on placebo analgesia in this study, we cannot rule out the possibility that OXT effects vary with different types of placebo effects.

An interesting nuance in our findings is that using a frequentist statistical linear mixed-effects model we found that OXT administration did statistically significantly increase the effect of branding efforts of an energy drink on cognitive performance after its consumption versus consumption of an identical soft drink. Participants were unconscious of this increase in performance. Both their reported anticipated performance and confidence in their performance stayed unchanged, which also might explain the absence of an interpretable main effect of branding. Our additional explorative Bayesian analysis, however, did not provide any robust evidence in favour of this interaction of OXT treatment with branding efforts. We call for future research that investigates the possibility that (1) the difference between evaluating one’s subjective enjoyment versus performing a numerical Stroop task and/or (2) the way we set up the branding manipulation did change the impact that OXT might have that is not directly linked to trust in the marketer and his branding efforts.

Regarding the difference in the type of tasks, it is noteworthy that another study investigating the role of OXT for expectancy effects on working memory performance — a different type of cognitive performance — found that OXT administration did significantly increase expectancy-driven performance^52,53^. Thus, future research needs to address whether OXT might play a different role when evaluating one’s subjective experience versus performing a cognitive task.

One difference of relevance in how we set up the branding versus price/organic label task was that the branding task required more social interaction between the experimenter and participant. More specifically, the experimenter gave information about the effectiveness of the energy drink versus neutral information. Previous research has demonstrated that such social interactions are important for OXT to have behavioural effects, especially in the context of expressing conformity with trustworthy people who might be seen as experts^54–57^. Because we did not include any measures that could provide empirical proof that OXT did increase how trustworthy or knowledgeable the experimenter was perceived to be and that this might have led to increases in the impact of OXT on MPE, these ideas remain to be tested in future research.

Another limitation of our study is that we included only male participants. There is accumulating evidence that effects of intranasal OXT differ substantially between women and men^58^, which may be related to interactions between OXT and sex hormones such as estradiol^59^. We focused on an exclusively male sample to increase the statistical power and to control for an additional source of noise due to potential gender-related (hormonal) differences of OXT effects on behaviour^58,60,61^. However, focusing on male participants prevents us from making any reliable conclusion about the impact of OXT and trust on MPE in females.

It could also be seen as a limitation that we did not asses our participants’ blood- or saliva-levels of OXT to show that the nasal OXT administration worked. However, it is common practice in the OXT research to rely on previous studies demonstrating that intranasally administered OXT increases endogenous OXT levels and reaches the brain^62–65^. Moreover, the reliability and comparability of OXT measurements after intranasal administration is currently debated: Saliva concentrations might be biased as intranasally administered OXT can reach the saliva via the pharynx. Thus, saliva concentrations also reflect non-absorbed exogenous OXT. Measured concentrations in the blood vary strongly depending on the exact method of sample preparation and assay used^66^.

To conclude, our results suggest that neither OXT administration nor reported trust in various marketing actions statistically significantly increases MPE across those marketing actions and across different tasks (i.e., judging taste pleasantness and performing a cognitive task). Our findings make an important contribution to psychobiological theories of placebo effects across domains. They also provide marketers new insights into the role trust and social cognitions play in consumers’ reactions to marketing activities and how that affects decision-making and satisfaction.

## Materials and Methods

### Ethical Considerations and Open Science Reporting

The ethics committee of the German university approved this study. All participants gave written informed consent according to the Declaration of Helsinki and received monetary compensation for their participation. The study design and analyses plan were pre-registered at Open Science Framework (OSF) (https://osf.io/v3b2u, date of pre-registration: 7 March 2018) prior to the start of the data collection. A detailed description of any deviations from the pre-registration and the reasons for the deviations can be found in the supplementary material. Any exploratory analyses that we did not pre-register are labelled as such. Data were collected before the final data analysis presented in this paper started, and data is available on OSF (https://osf.io/9g4js/).

### Overview of experimental procedure

The experimental procedure is displayed in Figure 1a, and all details are provided in the supplementary material. The translated study materials are available on OSF (https://osf.io/9g4js/). Participants completed several questionnaires about their current mood, health, and general trust prior to the self-administration of the nasal spray that either contained OXT or was a sham spray. In line with previous studies^62,64^, intranasal OXT was administered 30 minutes before the start of the experiment. During the waiting time, participants watched a neutral animal documentary.

Afterwards, participants took part in a food tasting task followed by an exploratory cognitive performance task. Finally, we used questionnaires to sample their trust in the placebo cues used, their mood state after the tasks, belief about being in the treatment group, their socio-demographics, and the effectiveness of our marketing actions. The order of the tasks and questionnaires was identical for all participants. In total, the experiment lasted about 120 minutes. All tested participants received an average payment of €38.18, which was composed of a €30 baseline payment and up to €13 remuneration for their individual performance in the cognitive performance task. Debriefing about the exact aim of the study took place via email once the data collection was finished to avoid suspicion concerning our cover story due to communication between participants.

### Oxytocin administration

In a double-blind, between-group, sham-controlled design, participants intranasally self-administered either OXT (24 IU, Syntocinon® spray, Novartis, Basel, Switzerland; 6 puffs per nostril each with 2 IU) or a sham spray (6 puffs per nostril; identical appearance, smell, and ingredients except it did not contain the neuropeptide). The experimenter supervised the administration of the nasal spray, and the amount of administered spray was weighed. Additional puffs were administered when the administered amount fell below 600 mg. Dosage and administration procedure were based on the current literature about kinetics in men and guidelines for standardised nasal administration^62,67^.

### Materials and Methods Food Tasting Task

#### Participants

The overall sample size of this study was estimated based on a pilot study and is detailed in the OSF pre-registration, together with study inclusion criteria (https://osf.io/v3b2u). We tested 334 male participants aged between 18 and 59 years. We focused on male participants only for three reasons: 1) endogenous plasma levels of OXT have been shown to vary within women’s menstrual cycle^68^ and might confound experimental results and require us to control for menstrual cycle; 2) OXT effects can differ between men and women^58,60,61^, which adds additional noise to the data and requires much larger sample sizes; and 3) previous research has shown OXT effects on placebo analgesia in men^28^.

For the food tasting task, we applied two predefined exclusion criteria based on our pre-registration, which led to the exclusion of 109 participants from the analysis: 1) 97 participants were convinced that the products used in our study were identical, and 2) 16 participants recognized that the study was focused on MPE. (Four participants were excluded because both exclusion criteria applied to them.) We further excluded two participants based on two criteria that we did not foresee and did not pre-register but that were necessary to ensure data integrity: 1) 1 participant got a wrong dosage of nasal spray, and 2) 1 participant mixed up the order of the products and rated the wrong product. Thus, we excluded in total 111 datasets from analysis; the total sample size of the data for our main analysis was 223 (mean age ± SD: 26.70 ±7.94 years, *N*_sham_ = 111, mean age ± SD sham group: 27.33 ± 8.50 years; *N*_OXT_ = 112, mean age ± SD OXT group: 26.06 ± 7.31 years). The deviation from our pre-registered sample size is explained in the supplementary material and did not affect the results.

Participants were randomly assigned to either the OXT or sham group. Supplementary Table S13 shows the sample size and sociodemographic, personality, and other relevant characteristics of the sham and the OXT groups.

#### Stimuli

Participants tasted various chocolate truffles and 5-gram servings of applesauce. The product types, labels, and presentation (see Figure 1b) were chosen based on two pre-tests. First, we did an online pre-test asking participants (*N*=142) how good they would expect various foods with different product labels (presented as pictures) to taste. Second, we did a tasting experiment in the lab asking participants (*N*=25) to rate how much they liked the taste of different food products depending on labels and presentation. We chose chocolate truffles and applesauce because our labelling was able to elicit significant marketing placebo effects in these two product categories in the pre-tests (for details see supplementary material and study materials on OSF: https://osf.io/9g4js/).

#### Food tasting task

Each product type (i.e., chocolate truffles and applesauce) was paired with different marketing labels that have been shown to induce higher versus lower expectancies about how good the food should taste, as replicated in our pre-tests^3,4,18,69^. These were high versus low price tags for chocolate truffles and labels indicating whether the product was organic or not for the applesauce.

Participants received three chocolate truffles and three samples of applesauce, all labelled differently and presented to enhance how good the participants would expect the products to taste (Figure 1b). Note that when talking about label (effects) in our methods, results, and discussion sections we refer to the conjoint change in label and presentation that was pre-tested to enhance positive taste expectations. Unbeknownst to the participants, two of the three food samples were identical, but their labels and delivery differed (see supplementary Table S15 for the exact food products and other consumables of the food tasting task). The third sample was a different product from the same product category and served as a distractor so that participants could not easily see through the manipulation. Participants were instructed that the goal of this task was to test for the effects of OXT on taste perception.

We applied a two (treatment: OXT vs. sham, between-participants) by two (label and product packaging: positive and neutral, within-participant) mixed repeated-measures design in this task. Put differently, we did expect both types of labels (i.e., price tags and production standard labels) to act similarly. Hence, we pre-registered to treat the two label and product types as a repeated measure to increase the robustness of the design.

Participants were asked to taste each product and read its label carefully. After each tasting, participants rated how much they subjectively enjoyed the taste of the food on a 9-point Likert scale with anchors “not good at all” (coded as 1) and “exceptionally good” (coded as 9). The tasting was self-paced without any time limit. The order of the two product categories and the order of the three samples within each product category were randomized. After each sample, participants rinsed their mouths with water to neutralize their taste buds.

#### Data analysis food tasting task

All statistical tests for both tasks were conducted with the R language and environment^70^ (version 3.6.1), and we used the *lmer* function from the *robustlmm* package^71^ (version 2.3) for robust linear mixed-effects models. Based on recommendations for more robust and reliable model parameters, we decided on robust models as their estimates are less affected by violations of model assumptions^72,73^. The function for the robust linear mixed-effects model does not return confidence intervals and *p*-values. Therefore, 95% CIs were approximated by the Wald test with the *tab_model* function of the *sjPlot* package^74^ (version 2.8.6). Likewise, *p*-values were approximated by the Wald test assuming infinite degrees of freedom. All statistical tests were performed two-tailed with a significance threshold of α ≤ 0.05. All mixed-effects models with between- and within-participant variables contained random intercepts for participants to account for the repeated measures. We z-scored continuous variables across participants and across repeated measures. To test for the robustness of the obtained results, we repeated all models with covariates for demographic and psychometric control variables, as detailed in the supplementary materials.

We pooled the data of both product categories (i.e., chocolate truffles and applesauce) for all following analyses and analysed them as within-participant repeated measures unless otherwise stated.

We used a linear mixed-effects model to investigate whether we could successfully induce MPE on experienced taste pleasantness using positive labels. In this model, the dependent variable is the taste rating. Label (neutral = −0.5, positive = 0.5) and product type (chocolate truffles = −0.5, applesauce = 0.5) are used as fixed effects. In post hoc *t*-tests with Holm correction for multiple comparisons, we compared taste pleasantness ratings of positive and neutral labels separately for the two product types.

We applied a different linear mixed-effects model to test whether OXT administration increased MPE (MPE score = taste pleasantness rating_positive label_ – taste pleasantness rating_neutral label_). We regressed the MPE score across both product types on treatment (sham = −0.5, OXT = 0.5), product type (truffles = −0.5, applesauce = 0.5), and their two-way interaction.

We conducted a Bayesian analysis to quantify the evidence in favour of our observed null effect of OXT treatment on taste MPE. We ran a Bayesian mixed-effects model via the *lmBF* function with its default settings (JZS priors with an *r* scale of 0.5) from the *BayesFactor* package (Version 0.9.12-4.2)^75^. The model paralleled the frequentist linear mixed-effects model. We calculated the Bayes factor by comparing the full model (with all predictors) to the null model (= full model without the predictor for treatment) to quantify the evidence in favour of an OXT effect on MPE (BF10). We assessed the sensitivity of the Bayesian analysis to different priors for the fixed-effect treatment by repeating the analysis with a JZS prior with *r* scales of 0.25, 0.707, and 1.0.

We regressed the MPE score across both product types on trust in marketers across expensive and organic products (z-scored across participants and products), product type (truffles = −0.5, applesauce = 0.5), and their two-way interaction to test whether reported trust in marketing actions would increase MPE.

We used MPE score as a dependent variable and regressed it on taste and quality expectations (z-scored across participants and product types) and their two-way interactions with product type (chocolate truffles = −0.5, applesauce = 0.5), to explore the relationship of MPE and trust with product expectations (i.e., taste and quality expectations). In two additional models, we used either taste or quality expectations across both product types as a dependent variable and trust in expensive or organic products (z-scored across participants and product types), product type (chocolate truffles = −0.5, applesauce = 0.5), and their two-way interaction as independent variables.

To analyse whether OXT administration affects trust in expensive products and organically labelled products, we regressed the reported trust across both product types on treatment (sham = −0.5, OXT = 0.5), product type (truffles = −0.5, applesauce = 0.5), and their two-way interaction.

### Materials and Methods Cognitive Performance Task

#### Participants

For the cognitive performance task, we applied three predefined exclusion criteria based on our pre-registration, which led to the exclusion of 129 participants from the analysis: 1) 16 participants were convinced that the drinks were identical, 2) 113 participants reported not feeling any effect of the energy drink, and 3) 17 participants recognized that we were interested in MPE. (Seventeen participants were excluded because two or three exclusion criteria applied to them.) For the detailed questions in the manipulation check see uploaded the study material on OSF (https://osf.io/9g4js/). We further excluded three participants based on criteria that we did not foresee and did not pre-register but that were necessary to ensure data integrity: 1) 1 participant got a wrong dosage of nasal spray, and 2) 2 participants were not able to perform the cognitive performance task because they did not understand the instructions.

Thus, we excluded in total 132 datasets from analysis; the total sample size of the data for our main analysis was 202 (mean age ± SD: 26.5 ± 7.4 years, *N*_sham_ = 99, mean age ± SD sham group = 26.6 ± 7.3 years; *N*_OXT_ = 103, mean age ± SD OXT group = 26.3 ± 7.4 years). The final sample size exceeded the pre-registered sample size by 2 participants due to logistic reasons. Participants were randomly assigned to either the OXT or sham group. Supplementary Table S14 shows the sample size, socio-demographic, personality, and other relevant characteristics of the two groups.

#### Stimuli and procedure

We administered the same soft drink twice, once in a can of Schweppes Lemon© (a citrus-flavoured carbonated lemonade) and once in a can of Red Bull Silver Edition© (a citrus-flavoured carbonated energy drink) (see Figure 1c). Participants were informed that Red Bull Silver (which was not available and was unknown in Germany at the time of the study) was a new strong and fast-acting energy drink that had been tested and demonstrated to enhance performance within 5 to 10 minutes after ingestion, and that had measurable positive effects on mental performance for roughly 15 to 20 minutes. We specified that the energizing ingredients of the drink were caffeine and taurine, which are quickly taken up into the blood and able to stimulate the central nervous system. In support of our cover story, a self-made poster with a Red Bull Silver advertisement and scientific information about its ingredients and their effects on mental performance was placed on the wall in front of the participants^76^ (see study material on OSF: https://osf.io/9g4js/).

Our cover study’s supposedly short timing of the energy drink effects was necessary because of the limited amount of time available for completing all parts of the study. Reliable OXT effects can be expected for only 90 minutes after nasal spray administration^62^. Hence, both tasks of the study — the food tasting task and the cognitive performance task — had to be finished in about 90 minutes.

We informed participants that the goal of this task was to investigate whether OXT affects the stimulating properties of energy and soft drinks during exam-like tasks. Participants performed the cognitive performance tasks twice, once after consuming the drink labelled as a soft drink and once after consuming the drink labelled as an energy drink. The order of the drinks was randomized and counterbalanced across the two treatment groups. Participants were informed about the order of the drinks, and an open can (either a soft drink can or an energy drink can, according to the drink order) was brought to the testing room. For the energy drink can, participants were instructed to pour the whole amount (250 ml) into a disposable cup and drink all of it. For the soft drink can (300 ml) participants were instructed to pour the drink up to a 250-milliliter mark on the cup in order to consume the same amount of drink for both conditions.

#### Numerical Stroop task

After a short waiting time for the drink to “take effect”, participants completed one run of a numerical Stroop task (see Figure 1c) that was a modified version of a validated task designed to measure incentive motivation of cognitive effort^77^. The task consisted of 48 trials; during each trial, participants first saw the maximum number of points they could earn (in 50% of the trials the maximum was one point, in 50% it was ten points, randomized). To ensure motivation, the total amount of points earned was converted into an additional monetary payoff (1 point = 0.025 €).

The second screen of each trial showed a ladder with ten steps, each depicting a number pair. Participants’ task was to indicate the numerically higher number with the left or right arrow key; each correct answer allowed them to climb up to the next step. They had 3,500 milliseconds to climb up as far as possible on the ten steps. Each successful step of the ladder yielded one-tenth of the total number of points at stake. Numbers within each pair differed in their numerical (0–9) and physical size. The difference in physical size was identical for all pairs on the screen. The numerical difference varied between 1 and 5, with two pairs of each in all trials. If a participant made an error, the arrow pointing at the current number pair was locked for one second.

We also altered the difficulty of the task in a randomized fashion. In 50% of the trials the numerically greater number was also physically greater for all ten number pairs (i.e., 100% congruent). In the other 50% of the trials the numerically greater number was physically smaller (i.e., 50% congruent), which caused a numerical Stroop effect requiring more attentional effort^78–80^. After each trial, participants received feedback about their total accumulated number of points.

Before the first trial of each run, we asked participants to indicate how many number pairs they expected to indicate correctly. We used this anticipated performance as a measure of their expectation given the drink they thought they had consumed. Participants answered the same question about their expectations every sixth trial as a measure of their confidence in their performance.

#### Data analysis numerical Stroop task

We used a robust linear mixed-effects model to analyse performance (number of steps climbed in every trial) as a dependent variable. Fixed effects included label (soft drink = −0.5, energy drink = 0.5), trial (1–48 for the first run and 49–96 for the second run, mean cantered), run (first run = −0.5, second run = 0.5), reward (1 point = −0.5, 10 points = 0.5), treatment (sham = - 0.5, OXT = 0.5), order of drinks (soft drink first = −0.5, energy drink first = 0.5), and difficulty (100% congruent = easy = −0.5, 50% congruent = difficult = 0.5) as well as two-way interactions of treatment by label, treatment by difficulty, and treatment by reward. The interaction of treatment by label represents our main analysis of interest and tests whether the sham group and the OXT group differ in how the energy drink label affects their cognitive performance. We added the interactions of treatment by difficulty and treatment by reward to the model to control for possible confounding effects of the OXT treatment on how rewards and task difficulty affect cognitive performance. We included random intercepts per participant in the model to account for between-participant heterogeneity.

We performed a Bayesian analysis to quantify the evidence in favour of an effect of OXT treatment on cognitive performance. We ran a Bayesian mixed-effects model via the *lmBF* function with its default settings (JZS prior with an *r* scale of 0.5) from the *BayesFactor* package (Version 0.9.12-4.2)^75^. We compared the full model (paralleling the frequentist linear mixed-effects model) to the null model without the interaction of Treatment x Label to quantify the evidence in favour of an impact of OXT on the label effect against the null hypothesis (BF10). We assessed the sensitivity of the Bayesian analysis to different priors for the fixed-effect treatment by repeating the analysis with a JZS prior with *r* scales of 0.25, 0.707, and 1.0.

In a linear model, we regressed average performance across all trials of the first run on label (soft drink = −0.5, energy drink = 0.5), trust in branding (z-scored across participants), anticipated performance of the first run (z-scored across participants), and the two-way interactions of label by trust and label by anticipated performance to test whether trust in branding and/or anticipated performance moderated the effect of drink label on cognitive performance. We used only the data from the first run because anticipated performance of the second run was biased by the actual experienced performance from the first run. Thus, anticipated performance in the second run is not comparable to that of the first run. We used a Pearson correlation to examine whether trust in branding was associated with anticipated performance in the first run.

We used two linear models and regressed anticipated performance and confidence in performance separately on label (soft drink = −0.5, energy drink = 0.5), treatment (sham = 0,5, OXT = 0.5), trust (z-scored across participants), and the two-way interactions of label by treatment and label by trust to explore whether anticipated performance/confidence in performance were affected by the drink label and whether the OXT treatment and/or trust in branding moderated a potential drink label effect.

We conducted an unpaired two-sided *t*-test to compare reported trust in branding between sham and OXT groups.

## Questionnaires

We used several questionnaires embedded in different questionnaire blocks at different points during the study (see Figure 1a). Details about all administered questionnaires can be found in the supplementary material and in the study material on OSF (https://osf.io/hbsxw).

Most important, we assessed trust in marketing actions (questionnaire blocks 2 and 3) with a translated and modified version of the brand trust scale^81^. For the exact modifications, see the supplementary material. In questionnaire block 2 we assessed trust in marketers of expensive and organic products (as related to the tasting task), and in block 3 we assessed trust in marketers of branded products (as related to the cognitive performance task).

To ensure that participants in the sham and OXT groups did not randomly differ in their general ability to trust others, we assessed general trust with a translated representation of the revised NEO Personality Inventory^82^ from the International Personality Item Pool^83,84^ (questionnaire block 1).

## Data and material availability

Data for all participants and analyses scripts are publicly available on OSF (https://osf.io/9g4js/). All translated questionnaires are also provided at OSF (https://osf.io/hbsxw).

## Acknowledgements

We thank Aenne Läufer, Tien Khuc, and Sarah Kempf for help with data collection; Dr Holger Gerhardt and Kersten Diers for statistical advice; and Dr Peter Trautner for help in programming the numerical Stroop task. This work was funded by INSEAD R&D funds awarded to HP and the Diet-Body-Brain grant (grant number: 01EA1809B) from the Federal Ministry of Education and Research Germany awarded to BW. The funders had no role in study design, data collection and analysis, decision to publish, or preparation of the manuscript.

## Author contributions

HP, LS, DS, and DSS designed the study. DSS acquired, analysed, and visualized the data with input from HP, LS, and DS. HP and DSS wrote the first draft of the manuscript, which was revised by LS, DS, BW, and RH. HP, BW, and RH obtained funding, provided resources, and supervised the project. All authors edited and approved the final version of the manuscript.

