## Supplementary Material for "Investigation of whether oxytocin and trust play a role in placebo effects of marketing actions"

### 1. Supplementary Methods

#### 1.1. Participant recruitment and inclusion criteria

Participants were recruited via texts and flyers spread online, via email, and on the campus of the Bonn university and the Bonn university clinic (see study material at OSF: https://osf.io/9g4js/). The goal of the study was described as investigating the effects of the hormone oxytocin (OXT) on taste perception and cognitive performance.

The following pre-registered (https://osf.io/v3b2u) inclusion criteria were applied: age older than 17 and younger than 61, no kidney insufficiency or cardiovascular diseases, no excessive smoking (i.e., more than five cigarettes per day), no excessive energy drink consumption (i.e., more than once per week; designed to avoid participants with excessive consumption who are used to energy drink tastes and effects and might see through our manipulation in the cognitive performance task), proficient knowledge of the German language, no intake of any (psychoactive) prescription medication (except antiallergenic agents), no food allergies to any of the stimuli, and no past or current psychological disorders. To reduce the risk that participants would see through our experimental manipulation, we included only participants who had not already participated in similar studies and who had no educational background in psychology. Participants were asked about these inclusion criteria during recruiting via an online survey. An experimenter assessed current or past psychological disorders via the Mini-International Neuropsychiatric Interview (MINI)[1] prior to the start of the experiment. Based on the MINI, we could not enrol nine of the 344 invited participants in our study (due to indications of depression, generalized anxiety disorder, drug abuse, manic episodes, or antisocial personality disorder). One other invited participant could not be enrolled due to insufficient understanding of the German instructions. Thus, we tested in total 334 participants.

We also asked participants to commit to maintaining their regular bed and waking times, and to abstain from caffeine and any large meal 2 hours and energy drinks and alcohol 24 hours prior to the experiment. Adherence to these rules was assessed in a questionnaire; no further physical checks were conducted.

The desired sample size of 200 participants in each task was determined a priori by a power analysis in G*Power[2]. The calculation was based on an unpublished pre-test for the food tasting part and a working paper[3] for the cognitive performance part, with similar designs. Details are provided in the pre-registration (https://osf.io/v3b2u).

##### 1.2. Pre-test for choice of stimuli in both tasks

For the choice of food stimuli in the food tasting task, we tested different food stimuli, labels, and their presentation or tasting in one online and two lab-based pre-tests.

First, in the online pre-tests (142 male participants recruited via Clickworker, mean age (SD): 27.2 (± 5.2) years, 1.50 € expense allowance), we used pictures of four product types: chocolate truffles, chocolate bars, apple juice, and crisps. For the labels we used information about country of origin, price, and manufacturing. Each product contained all three types of information. All product types were presented three times, with one of the three pieces of information (either price or manufacturing) varying across the three repetitions, while the other information and product appearance were kept identical. For the crisps, chocolate truffles, and chocolate bars we varied the price label and presented identical food pictures with a high (positive label), medium (distractor label), and low price tag (neutral label). For the apple juice, we varied the information about the manufacturing and presented identical pictures of apple juice with “organic” (positive label), “traditional” (distractor label), and “natural” (neutral label) description of manufacturing.

We randomized the order of the product types as well as the order of the three products within each product type, and participants rated their expected taste pleasantness of all presented products on a 9-point Likert scale.

We found that a high price tag (positive label) as compared to a low price tag (neutral label) statistically significantly increased expected taste pleasantness for all three food products, with the strongest effect for the chocolate truffles (paired, two-sided *t*-test, $p < 0.001$). The "organic" (positive) label did not statistically significantly affect expected taste pleasantness of the apple juice as compared to the "natural" (neutral) label (paired, two-sided *t*-test, $p > 0.216$).

We conducted an additional pre-test in our laboratory with a modified neutral label for the apple juice (using "naturally cloudy" instead of "natural," as the former might be better suited to induce lower expectations compared to the "organic" label). In addition to the apple juice, we also used applesauce as an alternative product with organic manipulation. For the price manipulation, we used chocolate bars and chocolate truffles. We used distinct packaging for each product (i.e., more vs. less exclusive packaging for the price manipulation and more vs. less natural packaging for the organic manipulation) to enhance the expectation effect of the label. We recruited 25 male and female participants in this paper-and-pencil version of the task. Products were provided by the experimenter and tasted in non-randomized fashion. We found statistically significant effects on the experienced taste pleasantness ratings for applesauce ($F_{1,24} = 5.616$, $p = 0.0262$) and chocolate truffles ($F_{1,24} = 7.111$, $p = 0.0135$).

We conducted a pre-test with 17 male and female participants in our laboratory to test which soft drink (choice set: Schweppes Lemon®, Sprite®, Campari Group LemonSoda®, San Pellegrino Limonata®) was somewhat but not too close in taste to Red Bull Silver Edition® and would be perceived as energizing. This was important so that participants would not see through our cover story. In the first part of the blind tasting, participants rated each soft drink for its taste pleasantness, how refreshing and how energizing they found it, and how familiar the drink was to them. In the second part of the blind tasting, participants received the same four soft drinks but were informed that one of them would be the Red Bull Silver Edition® and were asked to guess which one. Overall, the pre-test revealed that Schweppes Lemon® resembled an energy drink most as it was least familiar (tasted familiar to 11% of the participants) and tasted most similar to an energy drink (declared to be the energy drink by 41% of the participants).

##### 1.3. Additional information for the experimental procedure

Participants arrived in the lab between 8:30 am and 5:15 pm and verbally confirmed that they had not taken any medication since the online screening and were not suffering from nasal congestion (for a successful nasal spray administration). Only one participant was invited per session to avoid distraction and interaction between participants, which might have compromised our manipulation. There was always one experimenter (all four involved experimenters were female) present in the room while the participants conducted all questionnaires and tasks.

##### 1.4. Additional information on questionnaires

We assessed trust in marketing actions with a custom-translated and modified version of the brand trust scale[4]. Modifications of the original brand trust scale consisted of using specific marketing labels ("expensive products", "organic products", "brand products") instead of the placeholder "brand [X]", employing a 7-point Likert scale instead of a 5-point Likert scale, omitting one intentionality item ("I could rely on [X] brand to solve the problem."), and adding one reliability item ("With the purchase of brand [X], I get what I expect from a product."). We further added two questions asking about taste and quality expectations of expensive/organic/brand products to the questionnaire.

We measured autistic-like personality traits with the Autism Spectrum Quotient[5] (AQ; Figure 1a, questionnaire block 1). This served two purposes: first, we wanted to make sure that participants of the sham and OXT group did not differ in their autistic-like personality traits, as (a) they might affect social and non-social behaviors[6–8] and (b) they might affect the effect that OXT administration has on behaviors[9,10]. Second, we wanted to test directly whether autistic-like personality traits moderated OXT effects on our dependent variables, since previous studies found such effects for brand relationships[9].

We sampled positive and negative affect (PANAS[11]) in questionnaire block 1 (Figure 1a) before OXT administration and in questionnaire block 3 (Figure 1a) after the experiment to control for any potential confounding effects of OXT on mood.

Questionnaire block 1 (Figure 1 a) further contained several control questions about current health and well-being; questions about current disease symptoms were included to rule out the possibility that physical discomfort would affect behaviour in our tasks. The reported extent of physical activity on the study day was checked because extensive physical activity could bias endogenous OXT levels. No participant mentioned any current health issue or major sports activity that would interfere with our OXT treatment. Moreover, we sampled current hunger level and sleep (duration during the previous night and time since waking up) as control variables for the tasting and cognitive performance tasks, respectively.

##### 1.5. Waiting time with filler task

During the 30 minutes of waiting time after nasal spray administration, participants watched an animal documentary to keep their activity as comparable as possible. The documentary lasted 20 minutes. Two control questions about participants' happiness during the movie and the emotionality of the movie ensured that the moods of the sham and OXT groups were not influenced differently (see Supplementary Table S13 and Supplementary Table S14).

In the remaining 10 minutes of waiting time, participants were verbally instructed about the upcoming tasks and the aims of the tasks. We explained that the purpose of the food tasting task was to test the impact of OXT on taste perception of different food products. For the cognitive performance task, we used a cover story to impact participants' beliefs about the drink and its effect on their performance. As part of this cover story we explained that the purpose of the cognitive performance task was to test the impact of OXT on cognitive performance and its interaction with the effects of caffeine and taurine. A translated version of the script used is included in the study materials (https://osf.io/9g4js/). Finally, participants read written instructions about the detailed procedure of the cognitive performance task before they started with the food tasting task.

##### 1.6. Decoy memory task

After the numerical Stroop task in each run, participants performed a memory task. This served as a decoy task to align with our cover story that we were testing energy drink effects on performance under typical academic pressure such as finance or accounting exams. In this task, participants had to memorize stock market symbols of 14 companies that were presented on the screen for 3 seconds each. This was followed by a waiting period of 30 seconds, during which participants were asked to write down as many European capitals or US states as possible. Afterwards, participants had 1 minute to identify seven stock market symbols. As pre-registered, the data for this task was not analysed because it served only as a decoy task.

##### 1.7. Changes in data analysis from pre-registered analysis plan

Data collection and study design took place according to our pre-registered plan. However, our data analysis differed in some points from the pre-registered analysis plan due to unexpected findings. All changes are described and explained below, and any additional exploratory analyses are labelled as such in the respective results or methods sections.

###### 1.7.1. Food tasting task

First, our sample size exceeded our pre-registered limit of 100 participants per treatment group. Our pre-registered sample size was calculated for the food tasting task. However, during data collection we had a higher participant exclusion rate for the cognitive performance task that led us to add more participants in total for both tasks than we pre-registered. We report the results of the larger sample size but verified that results regarding the pre-registered main hypotheses did not differ substantially for $N = 200$. That is, with a smaller sample size we also found statistically significant label effects on taste pleasantness (mean ± SD neutral labels: 6.13 ± 0.79, mean ± SD positive labels: 6.47 ± 0.79, $\beta = 0.34$, SE = 0.10, 95% CI [0.14, 0.55], $t = 3.32$, $p < 0.001$, $d = 0.43$) but no statistically significant OXT treatment effect on MPE (mean ± SD sham group: 0.28 ± 1.12, mean ± SD OXT group: 0.40 ± 1.13, $\beta = 0.11$, SE = 0.17, 95% CI [-0.21, 0.44], $t = 0.69$, $p = 0.49$, $d = 0.10$).

Second, given the lack of our pre-registered effects, we did not conduct a mediation analysis with trust in marketers as a putative mediator of OXT effects on MPE.

Third, we decided not to include random intercepts for the product type but rather included product type as a fixed effect and potential moderator. The two products with their different label types differed strongly in their baseline taste ratings as well as the strength of their label effects. Including the product type as additional moderator allowed us to control for potential ceiling or floor effects or differences in underlying psychological mechanisms. The results for our main hypothesis did not change due to this adjustment: That is, with a random intercept for product we also found statistically significant label effects on taste pleasantness ($\beta = 0.34$, SE = 0.10, 95% CI [0.15, 0.54], $t = 3.53$, $p < 0.001$, $d = 0.44$) but no statistically significant OXT treatment effect on MPE ($\beta = 0.09$, SE = 0.16, 95% CI [-0.22, 0.40], $t = 0.55$, $p = 0.583$, $d = 0.08$).

Fourth, we used effect coding (-0.5/0.5) for all of our binary variables for a more straightforward interpretation of the estimates. In general, for linear mixed-effects models, the coding of the variables does not change the statistical results but only the interpretation of the individual estimates and the intercept[12].

Fifth, given that against our pre-registered hypothesis we found null effects, we conducted Bayesian analyses to quantify the nature of our null effects. Quantifying the strength of a statistically significant effect or of a null result is especially relevant to rule out false positive or false negative findings due underpowered study designs[13,14]. Regarding a possible replication crisis in OXT research[15–18], testing for the validity of effects via Bayesian statistics is of particular importance.

Sixth, the planned secondary analysis exploring whether marketing placebo responders show stronger OXT treatment effects was not conducted. Due to the between-group comparisons for OXT effects, such an analysis would not be meaningful.

Seventh, we did not dichotomize the continuous variables general trust and AQ for the moderation analysis using its median because using continuous predictors and moderators in linear models is statistically superior to artificially dichotomizing them[19–21].

##### 1.7.2. Cognitive performance task

First, paralleling the analysis of the food tasting task, we used a robust model for the cognitive performance analysis to alleviate the impact of outliers on model estimates[22,23]. Using a robust model over a normal linear mixed-effects model did not change the results substantially. That is, with a non-robust model (*lmer* function from the *lmerTest* package, version 3.1.3[24]) we also found a statistically significant main effect of drink label across sham and OXT group on cognitive performance ($\beta$ = 0.06, SE = 0.02, 95% CI [0.02, 0.09], $t$ = 2.77, $p$ = 0.006) and a statistically significant interaction of OXT treatment with drink label on cognitive performance ($\beta$ = 0.10, SE = 0.04, 95% CI [0.02, 0.17], $t$ = 2.40, $p$ = 0.016,).

Second, we pre-registered that we would calculate the performance difference between the energy drink condition and the soft drink condition. This would have required us to calculate an average performance per drink condition and would not have allowed us to have predictors for the level of reward and difficulty, as these variables are specific for each trial. We instead decided to use cognitive performance of each trial as a dependent variable rather than average performance difference and included run and label as predictors in the model. This analysis of trial-level performance is based on the analysis of Schmidt et al.[3] and is more powerful because it also considers trial difficulty and reward. Moreover, we decided not to add a random slope per participant for the level of reward because the variance of this random effect was zero. Thus, this random effect could be dropped from the model without influencing the results. As a further modification of the model, we adjusted the variable coding and used effect coding (-0.5, 0.5) for all binary variables. In general, for linear mixed-effects models, the coding of the variables does not change the statistical results but only the interpretation of the individual estimates and the intercept[12].

Third, we complemented our frequentist analysis with a Bayesian analysis for the same reasons detailed above for the food tasting task.

Fourth, we adjusted our analysis of anticipated performance and confidence in performance. We realized that due to our within-participant design, reported anticipated performance and confidence in the second condition might be biased by the actual experienced performance in the first condition. Therefore, we decided to analyse only the anticipated performance and confidence ratings of the first condition with label as between-participant variables and without repeated measures for participants.

Fifth, the planned secondary analysis examining whether marketing placebo responders show stronger OXT treatment effects was not conducted for the reasons detailed above for the food tasting task.

Sixth, we did not conduct the pre-registered cost-benefit model-based analysis. This model-based analysis was aimed at more closely investigating a potential interactive effect of drink label and reward on cognitive performance, which we did not observe in our data.

#### 2. Supplementary Results

##### 2.1. Tests of randomization

The two groups (i.e., sham and OXT) of the two tasks (i.e., food tasting task and cognitive performance task) were compared separately via Wilcoxon rank-sum tests for all demographic and psychometric control variables (see Supplementary Table S13 and Supplementary Table S14).

By chance, the two treatment groups of the exploratory cognitive performance task differed statistically significantly in their consumption frequency of energy drinks (mean ± SD sham group: 2.69 ± 1.10, mean ± SD OXT group: 3.07 ± 1.21, $Z = -2.04$, $p = 0.041$). Apart from that, the two treatment groups of food tasting task and cognitive performance task did not statistically significantly differ in any of the above-mentioned variables.

##### 2.2. Measurement of mood

To control for potentially confounding effects of OXT on mood, participants filled in the Positive and Negative Affect Schedule (PANAS)[11] before treatment administration and after the experiment. For each study population (i.e., food tasting task and cognitive performance task), two robust linear mixed-effects models with either positive or negative affect as dependent variable and time (pre-experiment = -0.5, post-experiment = 0.5), treatment (sham = -0.5, OXT = 0.5), and the two-way interaction of time and treatment as fixed effects were estimated.

For the participants of the food tasting task, we found no main effects of treatment and time nor an interaction effect of treatment and time on positive mood (Supplementary Table S16). However, negative mood scores statistically significantly decreased over the course of the experiment (mean ± SD pre: 11.76 ± 2.39, mean ± SD post: 10.96 ± 1.54, $\beta = -0.55$, SE = 0.09,

95% CI [-0.72, -0.37], $t$ = -6.10, $p$ = < 0.001, $d$ = 0.34). Moreover, a statistically significant interaction of time and treatment ($\beta$ = -0.45, SE = 0.18, 95% CI [-0.80, -0.10] $t$ = -2.51, $p$ = 0.012, $d$ = 0.29) indicates that the OXT group had a stronger decrease in negative mood score from pre- to post-measurement than the sham group (see Supplementary Table S16). Neither positive nor negative mood differed statistically significantly between sham and OXT groups before the start of the experiment (see Supplementary Table S13).

The robust linear mixed-effects model analysis for positive and negative mood of the participants of the cognitive performance task revealed a statistically significant increase in positive mood scores (mean ± SD pre: 30.54 ± 3.23, mean ± SD post: 31.33 ± 3.24, $\beta$ = 0.79, SE = 0.32, 95% CI [0.16, 1.41], $t$ = 2.45, $p$ = 0.014, $d$ = 0.24) and a statistically significant decrease in negative mood scores from pre- to post-measurement (mean ± SD pre: 11.94 ± 1.81, mean ± SD post: 11.15 ± 1.81, $\beta$ = -0.47, SE = 0.11, 95% CI [-0.68, -0.26], $t$ = -4.43, $p$ = < 0.001, $d$ = 0.26; see Supplementary Table S17). Neither positive nor negative mood differed statistically significantly between sham and OXT groups before the start of the experiment (see Supplementary Table S14). To control for these changes in mood, we included the difference in positive and negative PANAS scores (post- minus pre-score) as covariates to our main regression models of the food tasting and cognitive performance tasks. All observed results were robust when controlling for mood, as indicated in model 2 of the respective supplementary tables.

##### 2.3. Belief about the treatment and adverse effects

After the experiment, participants were asked to guess which treatment they received (OXT, sham, or no idea). Blinding of the treatment was validated with chi-squared tests separately for participant populations of the two tasks (i.e., tasting and cognitive performance). Participants were unaware of the received treatment as the treatment guess of the OXT group did not differ statistically significantly from chance (participants included in food tasting task: correct guess = 29.6%, $\chi^2_{(2)}$ = 3.88, $p$ = 0.14; participants included in cognitive performance task: correct guess = 35.0%, $\chi^2_{(2)}$ = 2.58, $p$ = 0.27). Treatment groups also did not differ statistically significantly in their treatment guess (participants included in food tasting task: $\chi^2_{(2)}$ = 0.05, $p$ = 0.97; participants included in cognitive performance task: $\chi^2_{(2)}$ = 1.57, $p$ = 0.46). No participant reported any side effects.

##### 2.4. Oxytocin effects on product and taste valuation

In addition to the two identical but differently manipulated products, participants tasted and rated a third product that differed from the manipulated products and served as a distractor (see Figure 1b). Previous research in rodents and humans suggests that OXT affects eating behavior by decreasing the rewarding properties of (sweet-tasting) food[25–29]. The distractor product allowed us to rule out the possibility that OXT affected taste or product perception of our offered sweet food products independent of marketing labels. We ran a linear mixed-effects model with taste pleasantness ratings of the distractor products as a dependent variable and treatment (sham = -0.5, OXT = 0.5), product type (truffles = -0.5, applesauce = 0.5), and their two-way interaction as fixed effect, and added random intercepts for participants. The taste ratings across both distractor products were not statistically significantly influenced by the OXT treatment (mean ± SD in sham group: 6.53 ± 1.34, mean ± SD in OXT group: 6.55 ± 1.21, $\beta$ = 0.09, SE = 0.17, 95% CI [-0.25, 0.43], $t$ = 0.52, $p$ = 0.61, $d$ = 0.07; see Supplementary Table S18).

##### 2.5. Oxytocin effects on taste and quality expectations

A positive attitude towards products does not necessarily need to rely only on trust. Positive product beliefs and expectations are well-known factors that reflect consumer attitudes and influence preferences and behaviour[30–33]. We analysed whether OXT influenced label-induced taste and quality expectations. As part of the trust questionnaire participants indicated their taste and quality expectations concerning expensive and organic products on a 7-point Likert scale. With robust linear mixed-effects models including random intercepts for participants, we estimated the effect of OXT on taste and quality expectations separately. OXT treatment changed neither taste (mean ± SD sham group: 4.00 ± 0.99, mean ± SD OXT group: 3.82 ± 1.14, $\beta$ = -0.17, SE = 0.14, 9% CI [-0.46, 0.11], $t$ = -1.21, $p$ = 0.23, $d$ = 0.16) nor quality expectations (mean ± SD sham group: 4.73 ± 1.08, mean ± SD OXT group: 4.60 ± 1.05, $\beta$ = -0.16, SE = 0.13, 95% CI [-0.41, 0.10], $t$ = -1.19, $p$ = 0.24, $d$ = 0.15) in a statistically significant way across both product types (see Supplementary Figure S3 and Supplementary Table S19 andSupplementary Table S20). Controlling for personality and demographic differences did not change the null effect of the OXT treatment (see model 2 in Supplementary Table S19 andSupplementary Table S20).

##### 2.6. General trust and autistic-like traits as potential moderators of treatment effects

As we defined in our pre-registration, we sought to explore whether general trust scores and autistic-like traits act as moderators of potential OXT effects. Previous research revealed differential OXT effects on empathy and consumer–brand relationships depending on autistic-like personality traits[9,10]. Based on this research, we wanted to test for similar differential OXT effects and included general trust scores and AQ scores as moderators in our main analyses.

For the food tasting task, we found statistically significant main effects of general trust on trust in marketers ($\beta$ = 0.48, SE = 0.19, 95% CI [0.10, 0.85], $t$ = 2.49, $p$ = 0.013; Supplementary Table S12) and on quality expectations ($\beta$ = 0.15, SE = 0.08, 95% CI [0.003, 0.30], $t$ = 2.00, $p$ = 0.045; Supplementary Table S20). These main effects mean that participants with higher levels of general trust in other people also have higher trust in marketing actions and higher quality expectations about expensive/organic products. We did not find any statistically significant main effect of general trust or AQ scores on MPE scores (general trust: $\beta$ = 0.03, SE = 0.09, 95% CI [-0.15, 0.21], $t$ = 0.31, $p$ = 0.753; AQ score: $\beta$ = 0.04, SE = 0.09, 95% CI [-0.14, 0.22], $t$ = 0.42, $p$ = 0.672) or taste expectations (general trust: $\beta$ = 0.03, SE = 0.08, 95% CI [-0.13, 0.20], $t$ = 0.39, $p$ = 0.699; AQ score: $\beta$ = -0.02, SE = 0.09, 95% CI [-0.19, 0.15], $t$ = -0.22, $p$ = 0.824). Moreover, we did not find any statistically significant interaction effect of general trust or autistic-like personality traits with OXT treatment on trust in marketing actions (general trust x treatment: $\beta$ = -0.22, SE = 0.37, 95% CI [-0.96, 0.51], $t$ = -0.59, $p$ = 0.554; AQ score x treatment: $\beta$ = 0.18, SE = 0.39, 95% CI [-0.58, 0.95], $t$ = 0.48, $p$ = 0.634), MPE scores (general trust x treatment: $\beta$ = 0.08, SE = 0.18, 95% CI [-0.27, 0.43], $t$ = 0.45, $p$ = 0.650; AQ score x treatment: $\beta$ = -0.07, SE = 0.19, 95% CI [-0.44, 0.29], $t$ = -0.40, $p$ = 0.693), taste expectations (general trust x treatment: $\beta$ = -0.21, SE = 0.17, 95% CI [-0.53, 0.12], $t$ = -1.25, $p$ = 0.211; AQ score x treatment: $\beta$ = -0.004, SE = 0.17, 95% CI [-0.34, 0.33], $t$ = -0.02, $p$ = 0.983), or quality expectations (general trust x treatment: $\beta$ = -0.08, SE = 0.15, 95% CI [-0.36, 0.21], $t$ =-0.52, $p$ = 0.604; AQ score x treatment: $\beta$ = 0.06, SE = 0.15, 95% CI [-0.24, 0.36], $t$ = 0.39, $p$ = 0.700). For all mentioned results see Supplementary Table S3, Supplementary Table S12, Supplementary Table S19, and Supplementary Table S20. These results indicate that stronger autistic-like personality traits or higher general trust was not related to a participant's sensitivity to OXT administration concerning our dependent variables.

As summarized in Supplementary Table S2, for the cognitive performance task, we did not find any statistically significant main effect of general trust ($\beta$ = 0.09, SE = 0.08, 95% CI [-0.07, 0.25], $t$ = 1.06, $p$ = 0.290) or AQ score ($\beta$ = 0.03, SE = 0.08, 95% CI [-0.14, 0.19], $t$ = 0.34, $p$ = 0.732) on cognitive performance. Moreover, we did also not find any statistically significant interaction of general trust ($\beta$ = 0.06, SE = 0.05, 95% CI [-0.03, 0.15], $t$ = 1.23, $p$ = 0.219) or AQ score ($\beta$ = 0.06, SE = 0.05, 95% CI [-0.03, 0.16], $t$ = 1.26, $p$ = 0.208) with the interactive effect of OXT and label on cognitive performance (i.e., three-way interaction of general trust or AQ score with treatment x label). We did, however, find a statistically significant interaction of AQ score with OXT treatment ($\beta$ = 0.42, SE = 0.17, 95% CI [0.08, 0.75], $t$ = 2.46, $p$ = 0.014). This interaction demonstrates that in the OXT group participants with a higher AQ score performed better as compared to people in the sham group with high AQ score.

#### 3. Supplementary Tables

**Supplementary Table S1**

*Robust linear mixed-effects model for experienced taste pleasantness depending on label and product type*

| | **DV: Experienced Taste Pleasantness** | | | |
|---|---|---|---|---|
| **Explanatory variables** | **Estimate (95% CI)** | **SE** | ***t*-value** | ***p*-value** |
| Intercept | 6.42 (6.28 – 6.56) | 0.07 | 88.6 | **<0.001** |
| Label (positive vs. neutral) | 0.34 (0.15 – 0.53) | 0.10 | 3.49 | **<0.001** |
| Product (applesauce vs. truffles) | -1.07 (-1.26 – -0.88) | 0.10 | -10.95 | **<0.001** |
| Label*Product | -0.17 (-0.56 – 0.21) | 0.20 | -0.89 | 0.372 |
| **Random Effects** | | | | |
| $\sigma^2$ | 2.02 | | | |
| $\tau_{00\ ID}$ | 0.61 | | | |
| $N_{ID}$ | 223 | | | |
| Observations | 892 | | | |
| Marginal $R^2$ / Conditional $R^2$ | 0.108 / 0.314 | | | |

*Notes*. Label and product type are both within-participant factors and are coded -0.5 for neutral label and 0.5 for positive label and -0.5 for chocolates and 0.5 for applesauce, respectively. CI = confidence interval; DV = dependent variable; SE = standard error of the estimate.

**Supplementary Table S2**

*Robust linear mixed-effects model for cognitive performance in numerical Stroop task*

| | DV: Cognitive performance of each trial | | | | | | | |
|---|---|---|---|---|---|---|---|---|
| | Model 1 | | | | Model 2 | | | |
| **Explanatory variables** | **Estimate (95% CI)** | **SE** | ***t*-value** | ***p*-value** | **Estimate (95% CI)** | **SE** | ***t*-value** | ***p*-value** |
| Intercept | 6.18 (6.04 – 6.32) | 0.07 | 86.87 | **<0.001** | 6.17 (6.03 – 6.31) | 0.07 | 84.1 | **<0.001** |
| Treatment (OXT vs. sham) | 0.02 (-0.26 – 0.30) | 0.14 | 0.14 | 0.885 | 0.04 (-0.26 – 0.35) | 0.15 | 0.29 | 0.772 |
| Label (energy drink vs. soft drink) | 0.06 (0.02 – 0.10) | 0.02 | 3.02 | **0.003** | 0.06 (0.02 – 0.11) | 0.02 | 2.92 | **0.003** |
| Trial | -0.01 (-0.01 – -0.004) | 0.0007 | -7.99 | **<0.001** | -0.01 (-0.01 – -0.00) | 0.001 | -8.06 | **<0.001** |
| Run (second vs. first) | 0.97 (0.89 – 1.05) | 0.04 | 24.39 | **<0.001** | 0.97 (0.89 – 1.06) | 0.04 | 22.4 | **<0.001** |
| Reward (10 vs. 1) | 0.06 (0.02 – 0.10) | 0.02 | 3.15 | **0.002** | 0.07 (0.02 – 0.11) | 0.02 | 3.02 | **0.003** |
| Order (energy drink first vs. energy drink second) | 0.13 (-0.15 – 0.41) | 0.14 | 0.91 | 0.363 | 0.07 (-0.23 – 0.37) | 0.15 | 0.48 | 0.633 |
| Difficulty (difficult vs. easy) | -2.34 (-2.38 – -2.30) | 0.02 | -117.6 | **<0.001** | -2.36 (-2.40 – -2.32) | 0.02 | -108.9 | **<0.001** |
| Treatment* Label | 0.09 (0.02 – 0.17) | 0.04 | 2.38 | **0.017** | 0.11 (0.02 – 0.19) | 0.04 | 2.50 | **0.013** |
| Treatment* Reward | -0.04 (-0.12 – 0.04) | 0.04 | -1.05 | 0.293 | -0.02 (-0.10 – 0.07) | 0.04 | -0.44 | 0.662 |
| Treatment* Difficulty | -0.12 (-0.20 – -0.04) | 0.04 | -3.04 | **0.002** | -0.14 (-0.22 – -0.05) | 0.04 | -3.14 | **0.002** |
| AQ | | | | | 0.03 (-0.14 – 0.19) | 0.08 | 0.34 | 0.732 |
| General Trust | | | | | 0.09 (-0.07 – 0.25) | 0.08 | 1.06 | 0.290 |
| Positive Affect (post – pre) | | | | | -0.002 (-0.15 – 0.14) | 0.07 | -0.03 | 0.977 |
| Negative Affect (post – pre) | | | | | -0.06 (-0.21 – 0.09) | 0.08 | -0.79 | 0.431 |
| Age | | | | | -0.23 (-0.43 – -0.03) | 0.10 | -2.24 | **0.025** |
| Treatment Guess | | | | | 0.03 (-0.12 – 0.17) | 0.07 | 0.36 | 0.723 |
| Income | | | | | -0.07 (-0.26 – 0.12) | 0.10 | -0.73 | 0.468 |

| | | | | | |
|---|---|---|---|---|---|
| Hours of Sleep | | 0.08 (-0.07 – 0.23) | 0.08 | 1.07 | 0.284 |
| Time Since Waking Up | | 0.07 (-0.08 – 0.22) | 0.08 | 0.91 | 0.363 |
| Energy Drink Consume | | -0.10 (-0.26 – 0.07) | 0.08 | -1.15 | 0.250 |
| Belief in Energy Drink Effects | | 0.08 (-0.09 – 0.25) | 0.09 | 0.90 | 0.370 |
| Treatment* AQ | | 0.42 (0.08 – 0.75) | 0.17 | 2.46 | **0.014** |
| Label*AQ | | -0.03 (-0.07 – 0.02) | 0.02 | -1.03 | 0.303 |
| Treatment* General Trust | | 0.08 (-0.23 – 0.40) | 0.16 | 0.52 | 0.602 |
| Label* General Trust | | -0.02 (-0.07 – 0.03) | 0.02 | -0.83 | 0.406 |
| Treatment* Label*AQ | | 0.06 (-0.03 – 0.16) | 0.05 | 1.26 | 0.208 |
| Treatment* Label* General Trust | | 0.06 (-0.03 – 0.15) | 0.05 | 1.23 | 0.219 |
| **Random Effects** | | | | | |
| $\sigma^2$ | 1.83 | 1.8 | | | |
| $\tau_{00}$ | 0.95 $_{ID}$ | 0.82 $_{ID}$ | | | |
| N | 202 $_{ID}$ | 168 $_{ID}$ | | | |
| Observations | 19,392 | 16,128 [a] | | | |
| Marginal $R^2$ / Conditional $R^2$ | 0.352 / 0.574 | 0.387 / 0.579 | | | |

*Notes.* Binary predictors are coded -0.5 and 0.5. Variable trial is mean centred; all other non-binary predictors are z-scored. AQ = Autism Spectrum Quotient; CI = confidence interval; DV = dependent variable; OXT = oxytocin; SE = standard error of the estimate.

[a] missing data points for variable income due to the voluntary nature of this question.

**Supplementary Table S3**

*Robust linear mixed-effects models for the impact of OXT treatment on MPE scores*

| | DV: MPE Score | | | | | | | |
|---|---|---|---|---|---|---|---|---|
| | Model 1 | | | | Model 2 | | | |
| **Explanatory variables** | **Estimate (95% CI)** | **SE** | ***t*-value** | ***p*-value** | **Estimate (95% CI)** | **SE** | ***t*-value** | ***p*-value** |
| Intercept | 0.33 (0.18 – 0.49) | 0.08 | 4.21 | **<0.001** | 0.35 (0.18 – 0.52) | 0.09 | 4.00 | **<0.001** |
| Treatment (OXT vs. sham) | 0.09 (-0.22 – 0.40) | 0.16 | 0.55 | 0.582 | 0.24 (-0.10 – 0.58) | 0.18 | 1.37 | 0.170 |
| Product (applesauce vs. truffles) | -0.24 (-0.55 – 0.07) | 0.16 | -1.54 | 0.124 | -0.17 (-0.51 – 0.17) | 0.17 | -0.99 | 0.323 |
| Treatment* Product | -0.09 (-0.70 – 0.53) | 0.31 | -0.28 | 0.781 | 0.01 (-0.67 – 0.69) | 0.35 | 0.03 | 0.979 |
| Positive Affect (post – pre) | | | | | -0.06 (-0.23 – 0.12) | 0.09 | -0.63 | 0.529 |
| Negative Affect (post – pre) | | | | | -0.03 (-0.21 – 0.14) | 0.09 | -0.37 | 0.713 |
| Product Liking | | | | | 0.02 (-0.16 – 0.19) | 0.09 | 0.17 | 0.864 |
| Treatment Guess | | | | | 0.08 (-0.10 – 0.25) | 0.09 | 0.84 | 0.401 |
| Age | | | | | -0.16 (-0.43 – 0.12) | 0.14 | -1.11 | 0.266 |
| Income | | | | | -0.01 (-0.25 – 0.23) | 0.12 | -0.06 | 0.950 |
| AQ | | | | | 0.04 (-0.14 – 0.22) | 0.09 | 0.42 | 0.672 |
| General Trust | | | | | 0.03 (-0.15 – 0.21) | 0.09 | 0.31 | 0.753 |
| Treatment*AQ | | | | | -0.07 (-0.44 – 0.29) | 0.19 | -0.40 | 0.693 |
| Treatment* General Trust | | | | | 0.08 (-0.27 – 0.43) | 0.18 | 0.45 | 0.650 |
| **Random Effects** | | | | | | | | |
| $\sigma^2$ | 2.77 | | | | 2.69 | | | |
| $\tau_{00}$ | 0.00 $_{ID}$ | | | | 0.00 $_{ID}$ | | | |
| N | 223 $_{ID}$ | | | | 190 $_{ID}$ | | | |
| Observations | 446 | | | | 379 [a] | | | |
| Marginal $R^2$ / Conditional $R^2$ | 0.006 / NA | | | | 0.021 / 0.021 | | | |

*Notes*. Treatment as between-participant factor is coded -0.5 for the sham group and 0.5 for the OXT group. Product as within-participant factor is coded -0.5 for chocolates and 0.5 for applesauce. All non-binary predictors are z-scored. AQ = Autism Spectrum Quotient; CI = confidence interval; DV = dependent variable; OXT = oxytocin; SE = standard error of the estimate.

[a] missing data points for variable income due to the voluntary nature of this question.

**Supplementary Table S4**

*Sensitivity analysis of different priors for the treatment regressor in the Bayesian model of the tasting task (Bayesian model paralleling frequentist model in Supplementary Table S3)*

| ***r* scale of JZS prior** | 0.250 (thin) | 0.50 (medium) | 0.707 (wide) | 1.00 (ultrawide) |
|---|---|---|---|---|
| **BF10** | 0.033 | 0.018 | 0.013 | 0.009 |

Notes. BF = Bayes Factor; JZS = Jeffreys-Zellner-Siow.

**Supplementary Table S5**

*Sensitivity analysis of different priors for the treatment regressor in the Bayesian model of the cognitive performance task (Bayesian model paralleling frequentist model in Supplementary Table S2)*

| ***r* scale of JZS prior** | 0.250 (thin) | 0.50 (medium) | 0.707 (wide) | 1.00 (ultrawide) |
|---|---|---|---|---|
| **BF10** | 0.218 | 0.159 | 0.131 | 0.131 |

Notes. BF = Bayes Factor; JZS = Jeffreys-Zellner-Siow.

**Supplementary Table S6**

*Robust linear mixed-effects model for the relationship of trust in marketing actions and MPE scores*

| | DV: MPE Score | | | | | | | |
|---|---|---|---|---|---|---|---|---|
| | Model 1 | | | | Model 2 | | | |
| **Explanatory variables** | **Estimate (95% CI)** | **SE** | ***t*-value** | ***p*-value** | **Estimate (95% CI)** | **SE** | ***t*-value** | ***p*-value** |
| Intercept | 0.34 (0.18 – 0.51) | 0.08 | 4.15 | **<0.001** | 0.35 (0.18 – 0.53) | 0.09 | 3.95 | **<0.001** |
| Trust in Marketing Actions | 0.09 (-0.08 – 0.26) | 0.09 | 1.07 | 0.283 | 0.10 (-0.08 – 0.28) | 0.09 | 1.06 | 0.290 |
| Product (applesauce vs. truffles) | -0.25 (-0.57 – 0.08) | 0.17 | -1.49 | 0.135 | -0.22 (-0.56 – 0.13) | 0.18 | -1.22 | 0.222 |
| Trust*Product | 0.002 (-0.33 – 0.34) | 0.17 | 0.01 | 0.992 | 0.01 (-0.35 – 0.37) | 0.18 | 0.06 | 0.953 |
| Positive Affect (post – pre) | | | | | -0.07 (-0.25 – 0.11) | 0.09 | -0.78 | 0.437 |
| Negative Affect (post – pre) | | | | | -0.06 (-0.23 – 0.12) | 0.09 | -0.61 | 0.543 |
| Liking | | | | | -0.002 (-0.18 – 0.18) | 0.09 | -0.02 | 0.983 |
| Treatment Guess | | | | | 0.06 (-0.11 – 0.24) | 0.09 | 0.70 | 0.481 |
| Age | | | | | -0.15 (-0.42 – 0.13) | 0.14 | -1.02 | 0.306 |
| Income | | | | | -0.02 (-0.27 – 0.22) | 0.12 | -0.20 | 0.840 |
| AQ | | | | | 0.04 (-0.15 – 0.23) | 0.09 | 0.42 | 0.673 |
| General Trust | | | | | 0.02 (-0.17 – 0.20) | 0.09 | 0.19 | 0.850 |
| **Random Effects** | | | | | | | | |
| $\sigma^2$ | 2.86 | | | | 2.72 | | | |
| $\tau_{00}$ | 0.00 $_{ID}$ | | | | 0.00 $_{ID}$ | | | |
| N | 223 $_{ID}$ | | | | 190 $_{ID}$ | | | |
| Observations | 446 | | | | 379 [a] | | | |
| Marginal $R^2$ / Conditional $R^2$ | 0.007 / 0.007 | | | | 0.018 / 0.018 | | | |

*Notes.* Product as within-participant factor is coded -0.5 for chocolates and 0.5 for applesauce. All non-binary predictors are z-scored. AQ = Autism Spectrum Quotient; CI = confidence interval; DV = dependent variable.

[a] missing data points for variable income due to the voluntary nature of this question.

**Supplementary Table S7**

*Robust linear mixed-effects model for the relationship of label-induced quality and taste expectations and MPE scores*

| | DV: MPE Score | | | | | | | |
|---|---|---|---|---|---|---|---|---|
| | Model 1 | | | | Model 2 | | | |
| **Explanatory variables** | **Estimate (95% CI)** | **SE** | ***t*-value** | ***p*-value** | **Estimate (95% CI)** | **SE** | ***t*-value** | ***p*-value** |
| Intercept | 0.35 (0.19 – 0.51) | 0.08 | 4.19 | **<0.001** | 0.36 (0.19 – 0.54) | 0.09 | 4.03 | **<0.001** |
| Quality Expect. | -0.13 (-0.31 – 0.06) | 0.10 | -1.30 | 0.195 | -0.16 (-0.36 – 0.05) | 0.10 | -1.51 | 0.130 |
| Taste Expect. | 0.20 (0.01 – 0.39) | 0.10 | 2.05 | **0.041** | 0.20 (0.001 – 0.39) | 0.10 | 1.97 | **0.049** |
| Product (applesauce vs. truffles) | -0.17 (-0.50 – 0.15) | 0.17 | -1.03 | 0.302 | -0.13 (-0.48 – 0.22) | 0.18 | -0.73 | 0.468 |
| Quality Expect.*Product | -0.03 (-0.41 – 0.35) | 0.19 | -0.15 | 0.881 | -0.10 (-0.50 – 0.30) | 0.20 | -0.49 | 0.621 |
| Taste Expect.*Product | 0.05 (-0.32 – 0.43) | 0.19 | 0.27 | 0.785 | 0.14 (-0.25 – 0.54) | 0.20 | 0.72 | 0.473 |
| Positive Affect (post – pre) | | | | | -0.05 (-0.23 – 0.12) | 0.09 | -0.60 | 0.548 |
| Negative Affect (post – pre) | | | | | -0.04 (-0.22 – 0.14) | 0.09 | -0.46 | 0.648 |
| Liking | | | | | 0.02 (-0.16 – 0.20) | 0.09 | 0.17 | 0.862 |
| Treatment Guess | | | | | 0.07 (-0.10 – 0.25) | 0.09 | 0.79 | 0.428 |
| Age | | | | | -0.16 (-0.44 – 0.12) | 0.14 | -1.11 | 0.269 |
| Income | | | | | -0.02 (-0.27 – 0.22) | 0.12 | -0.18 | 0.858 |
| AQ | | | | | 0.02 (-0.17 – 0.20) | 0.09 | 0.18 | 0.856 |
| General Trust | | | | | 0.03 (-0.15 – 0.22) | 0.09 | 0.35 | 0.727 |
| **Random Effects** | | | | | | | | |
| $\sigma^2$ | 2.85 | | | | 2.71 | | | |
| $\tau_{00}$ | $0.00_{ID}$ | | | | $0.00_{ID}$ | | | |
| N | $223_{ID}$ | | | | $190_{ID}$ | | | |
| Observations | 446 | | | | 379 [a] | | | |
| Marginal $R^2$ / Conditional $R^2$ | 0.015 / 0.015 | | | | 0.028 / 0.028 | | | |

*Notes.* Product as within-participant factor is coded -0.5 for chocolates and 0.5 for applesauce. All non-binary predictors are z-scored. AQ = Autism Spectrum Quotient; CI = confidence interval; DV = dependent variable; Expect. = Expectation.

[a] missing data points for variable income due to the voluntary nature of this question.

**Supplementary Table S8**

*Robust linear mixed-models for the relationship of trust in marketing actions and taste and quality expectations*

| | DV: Taste Expectation | | | | DV: Quality Expectation | | | |
|---|---|---|---|---|---|---|---|---|
| **Explanatory variables** | **Estimate (95% CI)** | **SE** | ***t*-value** | ***p*-value** | **Estimate (95% CI)** | **SE** | ***t*-value** | ***p*-value** |
| Intercept | 3.91 (3.79 – 4.03) | 0.06 | 62.88 | **<0.001** | 4.73 (4.63 – 4.83) | 0.05 | 93.01 | **<0.001** |
| Trust in Marketing Actions | 0.70 (0.57 – 0.83) | 0.06 | 10.85 | **<0.001** | 0.73 (0.63 – 0.83) | 0.05 | 13.85 | **<0.001** |
| Product (applesauce vs. truffles) | -0.33 (-0.58 – -0.09) | 0.12 | -2.69 | **0.007** | 0.17 (-0.03 – 0.37) | 0.10 | 1.70 | 0.089 |
| Trust in Marketing Actions*Product | 0.14 (-0.11 – 0.39) | 0.13 | 1.10 | 0.270 | -0.20 (-0.41 – 0.01) | 0.11 | -1.89 | 0.059 |
| **Random Effects** | | | | | | | | |
| $\sigma^2$ | 1.62 | | | | 1.08 | | | |
| $\tau_{00}$ | 0.00 $_{ID}$ | | | | 0.00 $_{ID}$ | | | |
| N | 223 $_{ID}$ | | | | 223 $_{ID}$ | | | |
| Observations | 446 | | | | 446 | | | |
| Marginal $R^2$ / Conditional $R^2$ | 0.244 / 0.244 | | | | 0.328 / 0.328 | | | |

*Notes.* Product as within-participant factor is coded -0.5 for chocolates and 0.5 for applesauce. All non-binary predictors are z-scored. CI = confidence interval; DV = dependent variable.

**Supplementary Table S9**

*Linear model testing the moderating effect of trust in marketing actions and anticipated performance (i.e., expectations) for the MPE on average cognitive performance in the first run of the numerical Stroop task*

| | DV: Average Cognitive Performance in First Run (label between-participants) | | | | | | | |
|---|---|---|---|---|---|---|---|---|
| | **Model 1** | | | | **Model 2** | | | |
| **Explanatory variables** | **Estimate (95% CI)** | **SE** | ***t*-value** | ***p*-value** | **Estimate (95% CI)** | **SE** | ***t*-value** | ***p*-value** |
| Intercept | 5.75 (5.62 – 5.88) | 0.07 | 88.29 | **<0.001** | 5.78 (5.64 – 5.92) | 0.07 | 82.5 | **<0.001** |
| Label (energy drink vs. soft drink) | 0.17 (-0.08 – 0.43) | 0.13 | 1.34 | 0.182 | 0.10 (-0.19 – 0.39) | 0.15 | 0.70 | 0.485 |
| Trust in Marketing Actions | -0.03 (-0.17 – 0.10) | 0.07 | -0.52 | 0.604 | -0.02 (-0.17 – 0.13) | 0.08 | -0.24 | 0.814 |
| Anticipated Performance | 0.20 (0.07 – 0.33) | 0.07 | 3.05 | **0.003** | 0.16 (0.01 – 0.31) | 0.08 | 2.08 | **0.039** |
| Label*Trust in Marketing Actions | 0.09 (-0.17 – 0.35) | 0.13 | 0.66 | 0.510 | 0.03 (-0.27 – 0.33) | 0.15 | 0.19 | 0.850 |
| Label*Anticipated Performance | -0.08 (-0.34 – 0.18) | 0.13 | -0.61 | 0.546 | -0.08 (-0.38 – 0.23) | 0.15 | -0.49 | 0.626 |
| Positive Affect (post – pre) | | | | | -0.02 (-0.16 – 0.13) | 0.07 | -0.22 | 0.827 |
| Negative Affect (post – pre) | | | | | -0.09 (-0.24 – 0.05) | 0.07 | -1.26 | 0.210 |
| General Trust | | | | | 0.11 (-0.05 – 0.27) | 0.08 | 1.40 | 0.163 |
| AQ | | | | | 0.09 (-0.07 – 0.24) | 0.08 | 1.14 | 0.254 |
| Age | | | | | -0.13 (-0.32 – 0.07) | 0.10 | -1.30 | 0.197 |
| Treatment Guess | | | | | 0.04 (-0.10 – 0.19) | 0.07 | 0.62 | 0.534 |
| Income | | | | | -0.17 (-0.34 – 0.01) | 0.09 | -1.86 | 0.065 |
| Hours of Sleep | | | | | 0.004 (-0.14 – 0.15) | 0.07 | 0.05 | 0.960 |
| Time since Waking Up | | | | | 0.07 (-0.08 – 0.21) | 0.07 | 0.94 | 0.351 |
| Energy Drink Consume | | | | | -0.03 (-0.19 – 0.13) | 0.08 | -0.38 | 0.706 |

| | | | | | |
|---|---|---|---|---|---|
| Belief in Energy Drink Effects | | 0.09 (-0.07 – 0.25) | 0.0 8 | 1.09 | 0.279 |
| Observations | 202 | | 168 [a] | | |
| $R^2$ / $R^2$ adjusted | 0.061 / 0.038 | | 0.155 / 0.066 | | |

*Notes.* We measured anticipated performance in the form of expectations about performance before the start of the task. We used only the data of the first run for the analysis because anticipated performance of the second run could be biased by the experienced performance in the first run. The binary variable label is coded -0.5 for soft drink and 0.5 for energy drink. All other non-binary variables are z-scored. AQ = Autism Spectrum Quotient; CI = confidence interval; DV = dependent variable; OXT = oxytocin; SE = standard error of the estimate.

[a] missing data points for variable income due to the voluntary nature of this question.

**Supplementary Table S10**

*Linear model for testing the interactive effects of label and treatment and label and trust in branding on anticipated performance (i.e., expectations) in the first run of the numerical Stroop task*

| | **DV: Anticipated Performance in First Run (label between-participants)** | | | | | | | |
|---|---|---|---|---|---|---|---|---|
| | **Model 1** | | | | **Model 2** | | | |
| **Explanatory variables** | **Estimate (95% CI)** | **SE** | ***t*-value** | ***p*-value** | **Estimate (95% CI)** | **SE** | ***t*-value** | ***p*-value** |
| Intercept | 5.49 (5.19 – 5.80) | 0.15 | 35.54 | **<0.001** | 5.54 (5.21 – 5.86) | 0.16 | 33.8 | **<0.001** |
| Label (energy drink vs. soft drink) | 0.30 (-0.31 – 0.91) | 0.31 | 0.98 | 0.331 | 0.48 (-0.19 – 1.15) | 0.34 | 1.43 | 0.155 |
| Trust in Branding | -0.08 (-0.39 – 0.23) | 0.16 | -0.50 | 0.621 | -0.22 (-0.58 – 0.13) | 0.18 | -1.25 | 0.215 |
| Treatment (OXT vs. sham) | -0.32 (-0.93 – 0.29) | 0.31 | -1.03 | 0.304 | -0.35 (-1.03 – 0.33) | 0.35 | -1.01 | 0.314 |
| Label*Trust in Branding | -0.12 (-0.74 – 0.51) | 0.32 | -0.37 | 0.709 | -0.31 (-1.02 – 0.39) | 0.36 | -0.88 | 0.382 |
| Label*Treatment | 0.20 (-1.02 – 1.43) | 0.62 | 0.33 | 0.741 | -0.02 (-1.37 – 1.33) | 0.68 | -0.03 | 0.978 |
| Positive Affect (post – pre) | | | | | 0.18 (-0.15 – 0.51) | 0.17 | 1.06 | 0.291 |
| Negative Affect (post – pre) | | | | | -0.18 (-0.53 – 0.16) | 0.17 | -1.07 | 0.288 |
| General Trust | | | | | 0.09 (-0.27 – 0.46) | 0.19 | 0.50 | 0.618 |
| AQ | | | | | -0.14 (-0.50 – 0.23) | 0.18 | -0.74 | 0.462 |
| Age | | | | | -0.24 (-0.69 – 0.22) | 0.23 | -1.03 | 0.305 |
| Treatment Guess | | | | | -0.10 (-0.43 – 0.23) | 0.17 | -0.59 | 0.556 |
| Income | | | | | 0.15 (-0.27 – 0.57) | 0.21 | 0.70 | 0.486 |
| Hours of Sleep | | | | | 0.06 (-0.27 – 0.40) | 0.17 | 0.37 | 0.714 |
| Time Since Waking Up | | | | | -0.12 (-0.46 – 0.21) | 0.17 | -0.72 | 0.474 |
| Energy Drink Consume | | | | | -0.01 (-0.39 – 0.36) | 0.19 | -0.08 | 0.940 |
| Belief in Energy Drink Effects | | | | | 0.09 (-0.28 – 0.47) | 0.19 | 0.48 | 0.628 |
| Observations | | 202 | | | | 168 | | |

| | | |
|---|---|---|
| $R^2$ / $R^2$ adjusted | 0.013 / -0.012 | 0.073 / -0.025 |

*Notes*. We measured anticipated performance in the form of expectations about performance before the start of the task. We used only the data of the first run for the analysis because anticipated performance of the second run could be biased by the experienced performance in the first run. The binary variable label is coded -0.5 for soft drink and 0.5 for energy drink. All other non-binary variables are z-scored. AQ = Autism Spectrum Quotient; CI = confidence interval; DV = dependent variable; OXT = oxytocin; SE = standard error of the estimate.

[a] missing data points for variable income due to the voluntary nature of this question.

**Supplementary Table S11**

*Linear model for testing the interactive effects of label and treatment and label and trust in branding on average confidence in performance in the first run of the numerical Stroop task.*

| | DV: Average Confidence in Performance in First Run (label between-participants) | | | | | | | |
|---|---|---|---|---|---|---|---|---|
| | Model 1 | | | | Model 2 | | | |
| **Explanatory variables** | **Estimate (95% CI)** | **SE** | ***t*-value** | ***p*-value** | **Estimate (95% CI)** | **SE** | ***t*-value** | ***p*-value** |
| Intercept | 4.40 (4.21 – 4.59) | 0.09 | 46.63 | **<0.001** | 4.46 (4.26 – 4.65) | 0.10 | 44.76 | **<0.001** |
| Label (energy drink vs. soft drink) | 0.16 (-0.22 – 0.53) | 0.19 | 0.82 | 0.411 | 0.09 (-0.31 – 0.50) | 0.21 | 0.45 | 0.655 |
| Trust in Branding | -0.03 (-0.22 – 0.16) | 0.10 | -0.27 | 0.785 | -0.05 (-0.27 – 0.16) | 0.11 | -0.48 | 0.630 |
| Treatment (OXT vs. sham) | 0.04 (-0.33 – 0.41) | 0.19 | 0.22 | 0.827 | 0.07 (-0.34 – 0.48) | 0.21 | 0.33 | 0.738 |
| Label*Trust in Branding | -0.04 (-0.42 – 0.34) | 0.19 | -0.21 | 0.836 | -0.13 (-0.56 – 0.30) | 0.22 | -0.61 | 0.545 |
| Label* Treatment | -0.28 (-1.03 – 0.46) | 0.38 | -0.75 | 0.453 | 0.02 (-0.80 – 0.84) | 0.42 | 0.05 | 0.963 |
| Positive Affect (post – pre) | | | | | 0.17 (-0.03 – 0.37) | 0.10 | 1.67 | 0.097 |
| Negative Affect (post – pre) | | | | | -0.11 (-0.32 – 0.10) | 0.11 | -1.03 | 0.305 |
| General Trust | | | | | 0.09 (-0.14 – 0.31) | 0.11 | 0.77 | 0.44 |
| AQ | | | | | -0.07 (-0.29 – 0.15) | 0.11 | -0.63 | 0.527 |
| Age | | | | | -0.32 (-0.60 – -0.04) | 0.14 | -2.29 | **0.024** |
| Treatment Guess | | | | | 0.02 (-0.18 – 0.22) | 0.10 | 0.24 | 0.810 |
| Income | | | | | -0.04 (-0.30 – 0.22) | 0.13 | -0.3 | 0.763 |
| Hours of Sleep | | | | | 0.10 (-0.10 – 0.30) | 0.10 | 0.97 | 0.331 |
| Time Since Waking Up | | | | | -0.08 (-0.29 – 0.12) | 0.10 | -0.79 | 0.433 |
| Energy Drink Consume | | | | | -0.09 (-0.31 – 0.14) | 0.12 | -0.76 | 0.450 |
| Belief in Energy Drink Effects | | | | | 0.09 (-0.14 – 0.32) | 0.12 | 0.76 | 0.449 |

| | | |
|---|---|---|
| Observations | 202 | 168 [a] |
| $R^2$ / $R^2$ adjusted | 0.007 / -0.018 | 0.131 / 0.039 |

*Notes.* We measured confidence in performance in the form of expectations about performance every sixth trial during the task. We used only the data of the first run for the analysis because anticipated performance of the second run could be biased by the experienced performance in the first run. The binary variable label is coded -0.5 for soft drink and 0.5 for energy drink. All other non-binary variables are z-scored. AQ = Autism Spectrum Quotient; CI = confidence interval; DV = dependent variable; OXT = oxytocin; SE = standard error of the estimate.

[a] missing data points for variable income due to the voluntary nature of this question.

**Supplementary Table S12**

*Robust linear mixed-effects models for trust in marketing actions*

| | DV: Trust in Marketing Actions | | | | | | | |
|---|---|---|---|---|---|---|---|---|
| | **Model 1** | | | | **Model 2** | | | |
| **Explanatory variables** | **Estimate (95% CI)** | **SE** | ***t*-value** | ***p*-value** | **Estimate (95% CI)** | **SE** | ***t*-value** | ***p*-value** |
| Intercept | 15.64 (15.31 – 15.96) | 0.17 | 93.75 | **<0.001** | 15.62 (15.26 – 15.98) | 0.18 | 85.26 | **<0.001** |
| Treatment (OXT vs. sham) | -0.05 (-0.71 – 0.60) | 0.33 | -0.16 | 0.876 | 0.09 (-0.62 – 0.81) | 0.37 | 0.25 | 0.799 |
| Product (applesauce vs. truffles) | 0.73 (0.32 – 1.14) | 0.21 | 3.50 | **<0.001** | 0.91 (0.46 – 1.36) | 0.23 | 3.99 | **<0.001** |
| Treatment*Product | -0.43 (-1.24 – 0.39) | 0.42 | -1.02 | 0.306 | 0.10 (-0.80 – 0.99) | 0.46 | 0.21 | 0.834 |
| Positive Affect (post – pre) | | | | | 0.55 (0.19 – 0.91) | 0.18 | 2.96 | **0.003** |
| Negative Affect (post – pre) | | | | | 0.20 (-0.17 – 0.57) | 0.19 | 1.06 | 0.290 |
| Treatment Guess | | | | | 0.03 (-0.34 – 0.39) | 0.19 | 0.14 | 0.892 |
| Age | | | | | 0.01 (-0.56 – 0.59) | 0.29 | 0.04 | 0.966 |
| Income | | | | | 0.06 (-0.45 – 0.56) | 0.26 | 0.22 | 0.823 |
| AQ | | | | | -0.35 (-0.74 – 0.03) | 0.20 | -1.80 | 0.072 |
| General Trust | | | | | 0.48 (0.10 – 0.85) | 0.19 | 2.49 | **0.013** |
| Treatment*AQ | | | | | 0.18 (-0.58 – 0.95) | 0.39 | 0.48 | 0.634 |
| Treatment*General Trust | | | | | -0.22 (-0.96 – 0.51) | 0.37 | -0.59 | 0.554 |
| **Random Effects** | | | | | | | | |
| $\sigma^2$ | 4.59 | | | | 4.72 | | | |
| $\tau_{00}$ | 3.60 $_{ID}$ | | | | 3.49 $_{ID}$ | | | |
| N | 223 $_{ID}$ | | | | 190 $_{ID}$ | | | |
| Observations | 446 | | | | 380 [a] | | | |
| Marginal $R^2$ / Conditional $R^2$ | 0.017 / 0.449 | | | | 0.101 / 0.484 | | | |

*Notes*. Treatment as between-participants factor is coded -0.5 for the sham group and 0.5 for the OXT group. Product as within-participant factor is coded -0.5 for chocolates and 0.5 for applesauce. All non-binary predictors are z-scored. AQ = Autism Spectrum Quotient; CI = confidence interval; DV = dependent variable; OXT = oxytocin; SE = standard error of the estimate.

[a] missing data points for variable income due to the voluntary nature of this question.

**Supplementary Table S13**

*Demographic, psychometric, and other control variables of the participant groups in food tasting task compared via Wilcoxon rank-sum tests*

| | **Sham N=111** mean (± SD) | **Oxytocin N=112** mean (± SD) | ***Z*** | ***p*-value** | **N** [a] |
|---|---|---|---|---|---|
| Age (years) | 27.33 (8.50) | 26.06 (7.31) | 1.01 | 0.314 | 223 |
| Income (€) | 666.58 (604) | 634.01 (586) | 1.16 | 0.247 | 192 |
| Years of education | 14.00 (2.22) | 14.08 (2.26) | -0.28 | 0.779 | 223 |
| AQ | 14.91 (4.65) | 15.52 (5.39) | -0.67 | 0.504 | 223 |
| General trust | 50.66 (6.99) | 49.56 (7.23) | 0.83 | 0. 405 | 223 |
| Hunger[1] | 3.00 (1.06) | 3.09 (1.09) | -0.64 | 0.525 | 223 |
| Liking chocolate truffles[2] | 4.84 (1.56) | 4.96 (1.50) | -0.48 | 0.632 | 222 |
| Liking applesauce[2] | 5.23 (1.40) | 5.21 (1.38) | 0.07 | 0.946 | 223 |
| Consumption frequency chocolate truffles[3] | 2.08 (0.79) | 2.14 (0.81) | -0.62 | 0.537 | 223 |
| Consumption frequency applesauce[3] | 2.97 (0.78) | 2.93 (0.76) | 0.35 | 0.725 | 223 |
| Movie – happiness[4] | 4.59 (0.89) | 4.62 (0.83) | -0.38 | 0.702 | 223 |
| Movie – emotionality[5] | 3.24 (1.42) | 3.19 (1.44) | 0.28 | 0.779 | 223 |
| PANAS – positive mood | 30.66 (5.27) | 30.59 (5.52) | 0.06 | 0.95 | 223 |
| PANAS – negative mood | 11.53 (1.97) | 11.99 (2.73) | -1.31 | 0.19 | 223 |

*Notes*. [1] 6-point Likert scale: 1 = not hungry at all, 6 = very hungry; [2] 7-point Likert scale: 1 = not at all, 7 = exceptionally good; [3] 8-point scale: 1 = never, 2 = < 1 x year, 3 = every few months, 4 = 1 – 3 x per month, 5 = 1–2 x per week, 6 = 3–4 x per week, 7 = 5–6 x per week, 8 = > 6 x per week; [4] 7-point Likert scale: 1 = very unhappy, 7 = very happy; [5] 7-point Likert scale: 1 = neutral, 7 =

very emotional. AQ = Autism Spectrum Quotient; PANAS = Positive and Negative Affect Schedule; SD = standard deviation.

[a] missing data points for variable income due to the voluntary nature of this question; missing data point for liking of chocolate truffles because one participant forgot to answer this question.

**Supplementary Table S14**

*Demographic, psychometric, and other control variables of the study population in cognitive performance task compared via Wilcoxon rank-sum tests*

| | **Sham N=99** mean (± SD) | **Oxytocin N=103** mean (± SD) | ***Z*** | ***p*-value** | **N** [a] |
|---|---|---|---|---|---|
| Age (years) | 26.61 (7.33) | 26.43 (7.42) | 0.23 | 0.817 | 202 |
| Income (€) | 709.84 (615.37) | 576.01 (519.55) | 1.82 | 0.069 | 170 |
| Years of education | 13.85 (2.34) | 13.91 (2.20) | -0.12 | 0.906 | 202 |
| AQ | 15.30 (4.42) | 15.94 (5.42) | -0.56 | 0.573 | 202 |
| General trust | 50.61 (7.81) | 49.07 (7.58) | 1.41 | 0.157 | 202 |
| Time since waking up (h) | 4.91 (2.81) | 4.98 (3.00) | 0.04 | 0.97 | 202 |
| Time of sleep (h) | 7.45 (1.04) | 7.15 (1.05) | 1.78 | 0.076 | 202 |
| Frequency drinking energy drinks[1] | 2.69 (1.10) | 3.07 (1.21) | -2.04 | **0.041** | 202 |
| Belief in energy drink effects[2] | 4.13 (1.13) | 4.34 (1.02) | -1.38 | 0.166 | 202 |
| Movie – happiness[3] | 4.75 (0.87) | 4.71 (0.72) | 0.2 | 0.841 | 202 |
| Movie – emotionality[4] | 3.17 (1.32) | 3.33 (1.41) | -0.96 | 0.337 | 202 |
| PANAS – positive mood | 30.41 (5.35) | 30.73 (4.99) | -0.58 | 0.562 | 202 |
| PANAS – negative mood | 11.53 (2.00) | 12.40 (3.63) | -1.66 | 0.096 | 202 |

*Notes*. [1] 8-point scale: 1 = never, 2 = < 1 x per year, 3 = every few months, 4 = 1–3 x per month, 5 = 1–2 x per week, 6 = 3–4 x per week, 7 = 5–6 x per week, 8 = > 6 x per week; [2] mean of three items: enhancement of focus and attention, enhancement of cognition, and enhancement of physical strengths; all three questions had a 7-point scale: 1 = strongly disagree, 7 = strongly agree; [3] 7-

point Likert scale: 1 = very unhappy, 7 = very happy; [4] 7-point Likert scale: 1 = neutral, 7 = very emotional. AQ = Autism Spectrum Quotient; PANAS = Positive and Negative Affect Schedule; SD = standard deviation.

[a] missing data points for variable income due to the voluntary nature of this question.

**Supplementary Table S15**

*Stimuli and consumables of food tasting task*

| Product | Type | Company |
|---|---|---|
| Chocolate truffles (positive and neutral product) | “Truffles fantaisie natural” | La Praline Gothenburg |
| Chocolate truffles (distractor product) | “Fine French Cocoa Truffles” | Chocolat Mathez |
| Applesauce (positive and neutral product) | “Apfelmus, aus ausgesuchten Äpfeln” | Gut&Günstig |
| Applesauce (distractor product) | “Apfelmus, extra Qualität” | HAK |
| Packaging chocolate truffles (positive product) | Wood, for 3 x 4 chocolates | Hussel |
| Packaging chocolate truffles (neutral product) | Plastic, for 2 x 6 chocolates, brown | Hussel |
| Packaging chocolate truffles (distractor product) | Cardboard, for 3 x 4 chocolates, creme-white (item number: 0530) | Wohlers Versandhandel |
| Packaging applesauce (positive product) | Palm leaf, small bowl, 80 ml, 8x8 cm, quadratic | Kaufdichgrün |
| Packaging applesauce (neutral product) | Cardboard bowl, 9x9x3 cm, angular | Papstar |
| Packaging applesauce (distractor product) | Fingerfood plates, PS, 7x7 cm, black | Papstar |

**Supplementary Table S16**

*Robust linear mixed-effects model results of positive and negative affect measurements of the study population in the food tasting task*

| | DV: Positive Affect | | | |
|---|---|---|---|---|
| **Explanatory variables** | **Estimate (95% CI)** | **SE** | ***t*-value** | ***p*-value** |
| Intercept | 30.76 (30.01 – 31.51) | 0.38 | 80.33 | **<0.001** |
| Treatment (OXT vs. sham) | 0.27 (-1.23 – 1.77) | 0.77 | 0.35 | 0.724 |
| Time (post vs. pre) | 0.26 (-0.33 – 0.86) | 0.31 | 0.87 | 0.386 |
| Treatment*Time | 0.42 (-0.78 – 1.62) | 0.61 | 0.69 | 0.493 |
| **Random Effects** | | | | |
| $\sigma^2$ | | 9.82 | | |
| $\tau_{00\ ID}$ | | 26.14 | | |
| $N_{ID}$ | | 223 | | |
| Observations | | 444 [a] | | |
| Marginal $R^2$ / Conditional $R^2$ | | 0.001 / 0.727 | | |
| | DV: Negative Affect | | | |
| **Explanatory variables** | **Estimate (95% CI)** | **SE** | ***t*-value** | ***p*-value** |
| Intercept | 11.04 (10.90 – 11.18) | 0.07 | 158.2 | **<0.001** |
| Treatment (OXT vs. sham) | 0.05 (-0.22 – 0.32) | 0.14 | 0.36 | 0.717 |
| Time (post vs. pre) | -0.55 (-0.72 – -0.37) | 0.09 | -6.10 | **<0.001** |
| Treatment*Time | -0.45 (-0.80 – -0.10) | 0.18 | -2.51 | **0.012** |
| **Random Effects** | | | | |
| $\sigma^2$ | | 0.84 | | |
| $\tau_{00\ ID}$ | | 0.61 | | |
| $N_{ID}$ | | 223 | | |
| Observations | | 444 [a] | | |
| Marginal $R^2$ / Conditional $R^2$ | | 0.057 / 0.452 | | |

*Notes*. Positive and negative affect were measured with the Positive and Negative Affect Schedule (PANAS). Time = pre- and post-experiment (coded -0.5 and 0.5, respectively). Treatment is coded -0.5 for sham and 0.5 for OXT. CI = confidence interval; DV = dependent variable; OXT = oxytocin; SE = standard error of the estimate.

[a] 2 missing values because of technical issues during questionnaire administration.

**Supplementary Table S17**

*Robust linear mixed-effects model results of positive and negative affect measurements of the study population in the cognitive performance task*

| | DV: Positive Affect | | | |
|---|---|---|---|---|
| **Explanatory variables** | **Estimate (95% CI)** | **SE** | ***t*-value** | ***p*-value** |
| Intercept | 31.1 (30.35 – 31.84) | 0.38 | 81.63 | **<0.001** |
| Treatment (OXT vs. sham) | 0.53 (-0.96 – 2.03) | 0.76 | 0.70 | 0.485 |
| Time (post vs. pre) | 0.79 (0.16 – 1.41) | 0.32 | 2.45 | **0.014** |
| Treatment*Time | 0.18 (-1.08 – 1.43) | 0.64 | 0.28 | 0.782 |
| **Random Effects** | | | | |
| $\sigma^2$ | | 9.76 | | |
| $\tau_{00\ ID}$ | | 22.94 | | |
| $N_{ID}$ | | 202 | | |
| Observations | | 402 [a] | | |
| Marginal $R^2$ / Conditional $R^2$ | | 0.007 / 0.704 | | |
| | **DV: Negative Affect** | | | |
| **Explanatory variables** | **Estimate (95% CI)** | **SE** | ***t*-value** | ***p*-value** |
| Intercept | 11.16 (11.00 – 11.32) | 0.08 | 136.57 | **<0.001** |
| Treatment (OXT vs. sham) | 0.18 (-0.14 – 0.50) | 0.16 | 1.07 | 0.283 |
| Time (post vs. pre) | -0.47 (-0.68 – -0.26) | 0.11 | -4.43 | **<0.001** |
| Treatment*Time | -0.42 (-0.83 – 0.001) | 0.21 | -1.96 | **0.05** |
| **Random Effects** | | | | |
| $\sigma^2$ | | 1.08 | | |
| $\tau_{00\ ID}$ | | 0.74 | | |
| $N_{ID}$ | | 202 | | |
| Observations | | 402 [a] | | |
| Marginal $R^2$ / Conditional $R^2$ | | 0.040 / 0.431 | | |

*Notes*. Positive and negative affect were measured with the Positive and Negative Affect Schedule (PANAS). Time = pre- and post-experiment (coded -0.5 and 0.5, respectively). Treatment is coded -0.5 for sham and 0.5 for OXT. CI = confidence interval; DV = dependent variable; OXT = oxytocin; SE = standard error of the estimate.

[a] 2 missing values because of technical issues during questionnaire administration.

**Supplementary Table S18**

*Robust linear mixed-effects model of experienced taste pleasantness of distractor products depending on treatment*

| | DV: Experienced Taste Pleasantness | | | |
|---|---|---|---|---|
| **Explanatory variables** | **Estimate (95% CI)** | **SE** | ***t*-value** | ***p*-value** |
| Intercept | 6.65 (6.48 – 6.82) | 0.09 | 76.57 | **<0.001** |
| Treatment (OXT vs. sham) | 0.09 (-0.25 – 0.43) | 0.17 | 0.52 | 0.605 |
| Product (applesauce vs. truffles) | -0.11 (-0.45 – 0.23) | 0.17 | -0.65 | 0.513 |
| Treatment*Product | 0.13 (-0.55 – 0.81) | 0.35 | 0.38 | 0.708 |
| **Random Effects** | | | | |
| $\sigma^2$ | 3.2 | | | |
| $\tau_{00\ ID}$ | 0 | | | |
| $N_{ID}$ | 223 | | | |
| Observations | 446 | | | |
| Marginal $R^2$ / Conditional $R^2$ | 0.002 / 0.002 | | | |

*Notes*. Treatment as between-participants factor is coded -0.5 for the sham group and 0.5 for the OXT group. Product as within-participant factor is coded -0.5 for chocolates and 0.5 for applesauce. CI = confidence interval; DV = dependent variable; OXT = oxytocin; SE = standard error of the estimate.

**Supplementary Table S19**

*Robust linear mixed-effects models for taste expectations*

| | DV: Taste Expectations | | | | | | | |
|---|---|---|---|---|---|---|---|---|
| | Model 1 | | | | Model 2 | | | |
| **Explanatory variables** | **Estimate (95% CI)** | **SE** | ***t*-value** | ***p*-value** | **Estimate (95% CI)** | **SE** | ***t*-value** | ***p*-value** |
| Intercept | 3.94 (3.80 – 4.08) | 0.07 | 54.9 | **<0.001** | 3.94 (3.78 – 4.10) | 0.08 | 48.71 | **<0.001** |
| Treatment (OXT vs. sham) | -0.17 (-0.46 – 0.11) | 0.14 | -1.21 | 0.226 | -0.17 (-0.49 – 0.14) | 0.16 | -1.07 | 0.286 |
| Product (applesauce vs. truffles) | -0.18 (-0.46 – 0.10) | 0.14 | -1.23 | 0.217 | -0.18 (-0.49 – 0.13) | 0.16 | -1.12 | 0.261 |
| Treatment* Product | -0.07 (-0.64 – 0.49) | 0.29 | -0.26 | 0.794 | 0.12 (-0.51 – 0.74) | 0.32 | 0.36 | 0.716 |
| Positive Affect (post – pre) | | | | | 0.15 (-0.01 – 0.31) | 0.08 | 1.81 | 0.070 |
| Negative Affect (post – pre) | | | | | 0.07 (-0.10 – 0.23) | 0.08 | 0.80 | 0.425 |
| Treatment Guess | | | | | -0.07 (-0.24 – 0.09) | 0.08 | -0.89 | 0.374 |
| Age | | | | | -0.09 (-0.34 – 0.17) | 0.13 | -0.69 | 0.493 |
| Income | | | | | -0.05 (-0.27 – 0.18) | 0.11 | -0.42 | 0.676 |
| AQ | | | | | -0.02 (-0.19 – 0.15) | 0.09 | -0.22 | 0.824 |
| General Trust | | | | | 0.03 (-0.13 – 0.20) | 0.08 | 0.39 | 0.699 |
| Treatment*AQ | | | | | -0.004 (-0.34 – 0.33) | 0.17 | -0.02 | 0.983 |
| Treatment* General Trust | | | | | -0.21 (-0.53 – 0.12) | 0.17 | -1.25 | 0.211 |
| **Random Effects** | | | | | | | | |
| $\sigma^2$ | 2.18 | | | | 2.28 | | | |
| $\tau_{00}$ | 0.00 ID | | | | 0.00 ID | | | |
| N | 223 ID | | | | 190 ID | | | |
| Observations | 446 | | | | 380 [a] | | | |
| Marginal $R^2$ / Conditional $R^2$ | 0.007 / 0.007 | | | | 0.028 / 0.028 | | | |

*Notes*. Treatment as between-participants factor is coded -0.5 for the sham group and 0.5 for the OXT group. Product as within-participant factor is coded -0.5 for chocolates and 0.5 for applesauce. All non-binary predictors are z-scored. AQ = Autism Spectrum Quotient; CI =

confidence interval; DV = dependent variable; OXT = oxytocin; SE = standard error of the estimate.

[a] missing data points for variable income due to the voluntary nature of this question.

**Supplementary Table S20**

*Robust linear mixed-effects models for quality expectations*

| | DV: Quality Expectations | | | | | | | |
|---|---|---|---|---|---|---|---|---|
| | Model 1 | | | | Model 2 | | | |
| **Explanatory variables** | **Estimate (95% CI)** | **SE** | ***t*-value** | ***p*-value** | **Estimate (95% CI)** | **SE** | ***t*-value** | ***p*-value** |
| Intercept | 4.77 (4.64 – 4.90) | 0.07 | 73.15 | **<0.001** | 4.77 (4.63 – 4.91) | 0.07 | 66.63 | **<0.001** |
| Treatment (OXT vs. sham) | -0.16 (-0.41 – 0.10) | 0.13 | -1.19 | 0.235 | -0.19 (-0.47 – 0.09) | 0.14 | -1.33 | 0.183 |
| Product (applesauce vs. truffles) | 0.39 (0.22 – 0.56) | 0.09 | 4.51 | **<0.001** | 0.40 (0.22 – 0.59) | 0.09 | 4.38 | **<0.001** |
| Treatment* Product | -0.17 (-0.51 – 0.17) | 0.17 | -0.99 | 0.324 | -0.15 (-0.52 – 0.21) | 0.18 | -0.83 | 0.409 |
| Positive Affect (post – pre) | | | | | 0.13 (-0.01 – 0.27) | 0.07 | 1.83 | 0.067 |
| Negative Affect (post – pre) | | | | | 0.06 (-0.09 – 0.20) | 0.07 | 0.76 | 0.448 |
| Treatment Guess | | | | | 0.04 (-0.11 – 0.18) | 0.07 | 0.49 | 0.627 |
| Age | | | | | -0.24 (-0.46 – -0.01) | 0.11 | -2.08 | **0.037** |
| Income | | | | | 0.13 (-0.07 – 0.33) | 0.10 | 1.29 | 0.197 |
| AQ | | | | | -0.08 (-0.23 – 0.08) | 0.08 | -0.98 | 0.328 |
| General Trust | | | | | 0.15 (0.003 – 0.30) | 0.08 | 2.00 | **0.045** |
| Treatment*AQ | | | | | 0.06 (-0.24 – 0.36) | 0.15 | 0.39 | 0.700 |
| Treatment* General Trust | | | | | -0.08 (-0.36 – 0.21) | 0.15 | -0.52 | 0.604 |
| **Random Effects** | | | | | | | | |
| $\sigma^2$ | 0.8 | | | | 0.77 | | | |
| $\tau_{00}$ | 0.50 ID | | | | 0.51 ID | | | |
| N | 223 ID | | | | 190 ID | | | |
| Observations | 446 | | | | 380 [a] | | | |
| Marginal $R^2$ / Conditional $R^2$ | 0.034 / 0.409 | | | | 0.090 / 0.450 | | | |

*Notes*. Treatment as between-participants factor is coded -0.5 for the sham group and 0.5 for the OXT group. Product as within-participant factor is coded -0.5 for chocolates and 0.5 for applesauce. All non-binary predictors are z-scored. AQ = Autism Spectrum Quotient; CI = confidence interval; DV = dependent variable; OXT = oxytocin; SE = standard error of the estimate.

[a] missing data points for variable income due to the voluntary nature of this question.

#### 4. Supplementary Figures

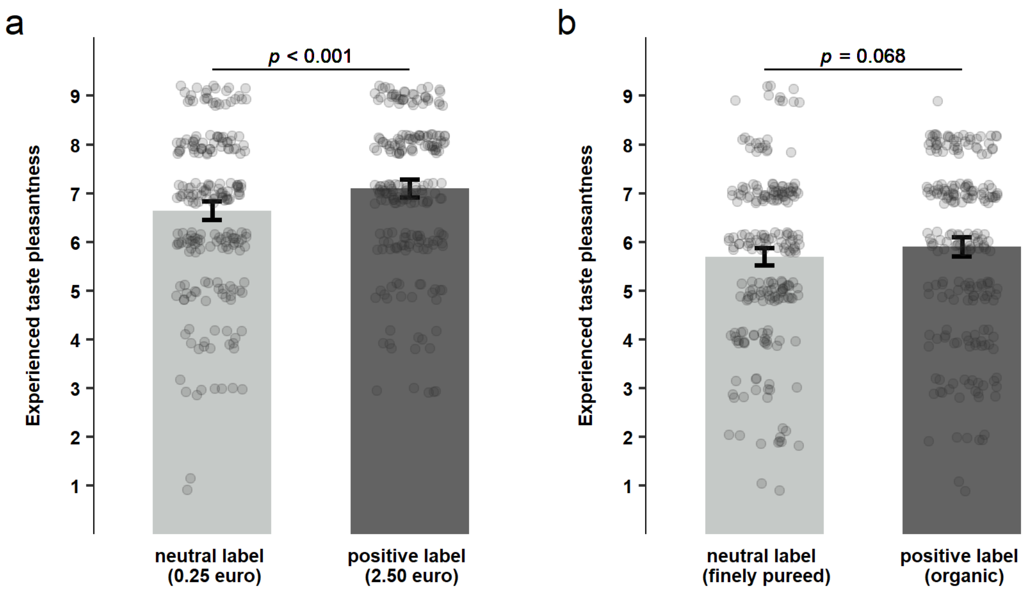

**Supplementary Figure S1: Effects of expensive and organic labels on experienced taste pleasantness of otherwise identical food products.** Experienced taste pleasantness compared for neutral and positive labels for expensive (a) and organic (b) labels separately. Individual observations ($N$ = 223 for each label across both treatment groups) are shown with grey dots. Error bars denote 95% confidence interval of the means. The expensive price tag had a significant positive impact on taste pleasantness ($t(222) = 4.18$, $p_{corr.} < 0.001$, $d = 0.28$, 95% CI of the difference [0.24, 0.67]) while the organic label did not (t(222) = 1.84, pcorr. = 0.068, d = 0.12, 95% CI of the difference [-0.02, 0.44]; paired, two-sided $t$-tests with Holm correction for $p$-values). For exact labels and appearance of products see Methods section and Figure 1b.

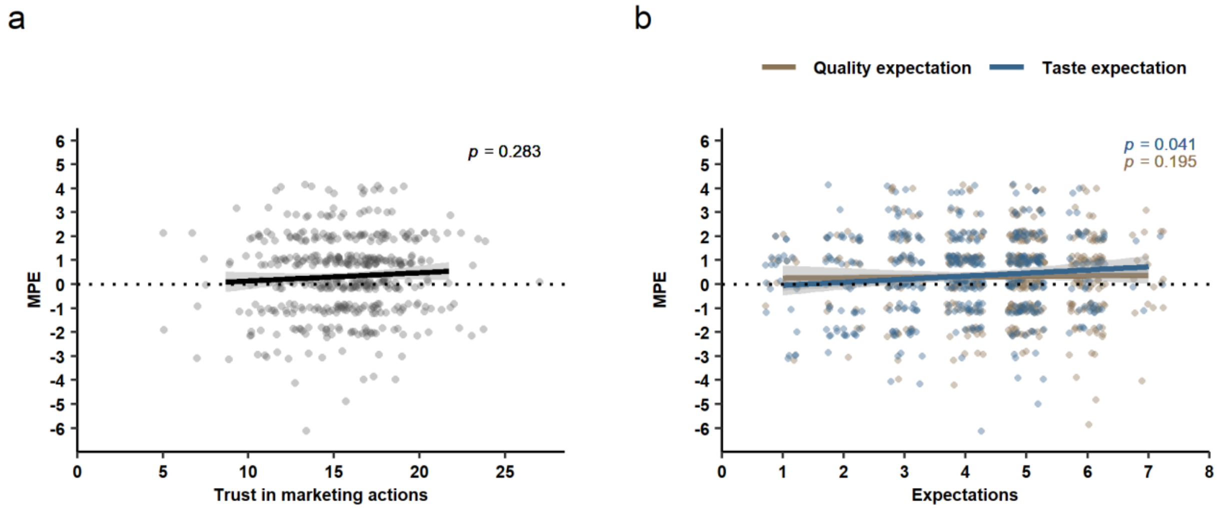

**Supplementary Figure S2: Analysis of the relation between marketing placebo effects and consumer trust and expectations.** (**a**) The relationship between trust in marketing actions ($\beta$ = 0.09, SE = 0.06, 95% CI [-0.08, 0.26], $t$ = 1.07, $p$ = 0.28) and (**b**) expectations (quality: $\beta$ = -0.13, SE = 0.10, 95% CI [-0.32, 0.06], $t$ = -1.30, $p$ = 0.195; taste: $\beta$ = 0.20, SE = 0.10, 95% CI [0.008, 0.39], $t$ = 2.05, $p$ = 0.041) with MPE. Dots represent individual data points of participants for both product types pooled across sham and OXT group ($N$ = 446 in (**a**) and $N$ = 446 per expectation type in (**b**)). Shaded area represents 95% confidence interval. MPE = marketing placebo effect.

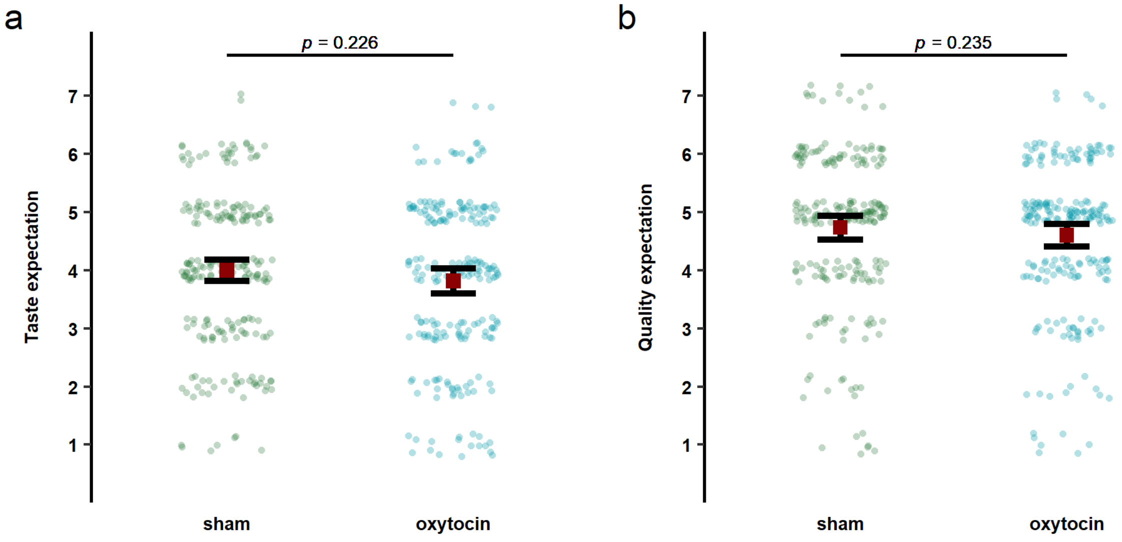

**Supplementary Figure S3: Evaluation of oxytocin's impact on taste and quality expectations.** (**a**) Comparison of taste expectations ($\beta$ = -0.17, SE = 0.14, 95% CI [-0.46, 0.11], $t$ = -1.21, $p$ = 0.23, $d$ = 0.16) and (**b**) quality expectations ($\beta$ = -0.16, SE = 0.13, 95% CI [-0.41, 0.10], $t$ = -1.19, $p$ = 0.24, $d$ = 0.15) between sham and OXT group. Dots represent individual data points of participants for both product types pooled ($N_{sham}$ = 222, $N_{OXT}$ = 224). We depicted mean values (red squares) with error bars (95% confidence intervals of the means) for ordinal Likert scale data.
